# Probabilistic mapping of sub-genic intolerance reveals functional and disease-critical protein regions

**DOI:** 10.64898/2026.09.13.745535

**Authors:** Constantine Stavrianidis, Yuncheng Duan, Grace E. Rhodes, Tristan J. Hayeck, William H. Majoros, Andrew S. Allen

## Abstract

Different regions of genes perform distinct functions and vary in their importance to human health. Evolutionary intolerance provides a powerful means of identifying regions where disruptive mutations are under strong purifying selection, informing genetic disease discovery and variant interpretation. However, estimating intolerance in small sub-genic regions from population variation alone is underpowered and unstable. We present PRIME, a Bayesian model that stabilizes estimates of regional missense intolerance by sharing information hierarchically across regions. Importantly, PRIME produces a full joint posterior across all genes, allowing complex inferential questions that are difficult or impossible to address with existing approaches to be answered. We utilize this to identify regions enriched for pathogenic and experimentally deleterious missense variants, improve prioritization of Mendelian disease genes by focusing on their most intolerant regions, and uncover conserved patterns of purifying selection across protein families. Integrating PRIME with existing computational variant predictors improves pathogenicity prediction, demonstrating that regional missense intolerance provides complementary information for clinical variant interpretation.

## Introduction

A key challenge in medical genetics is implicating variants in disease. Missense variants cause localized changes in encoded proteins through amino acid substitutions but do not always perturb function in the protein. Quantifying purifying selection against missense variants within a given gene or sub-genic region provides important information when predicting how damaging a variant found in such a region may be. This can be estimated by leveraging standing variation in the human population: genes that show a depletion of missense variants relative to expectation, referred to as intolerance, are likely subject to stronger purifying selection because variants in these genes are more likely to have disruptive effects on fitness and are therefore removed from the population over generations^1^. As the strength of purifying selection reflects functional importance, such *intolerant* genes are enriched for disease-causing variants^1–4^.

The first approach to measuring intolerance did so at the gene level^1^. Samocha et al.^2^ and Lek et al.^3^ proposed additional gene-level metrics that quantify variant depletion, which they referred to as genetic constraint. Throughout this paper, we will favor the antecedent term intolerance except when referring to existing constraint metrics. While useful for prioritizing potentially disease-causing genes, these methods do not capture intolerance that is localized to subregions within genes. Various parts of a protein may tolerate localized amino acid changes differently depending on structural or functional context. In fact, in some disease-associated genes, pathogenic missense variants are known to cluster within specific regions^5–13^.

Quantifying intolerance at a sub-genic level accounts for this intragenic heterogeneity and can help identify the functionally critical segments of a protein that are most sensitive to variation^14–20^. This finer resolution can improve variant interpretation, enhance gene-disease association studies, and provide deeper insights into how structural and functional features shape the distribution of evolutionary intolerance across proteins.

Current sub-genic methods differ in their approach to this problem. Previous work (subRVIS^14^) measures intolerance in subregions of genes defined by either protein domain or exon boundaries in a regression framework (RVIS^1^). Other methods define intolerant regions empirically, either by identifying exonic segments devoid of protein-altering variation (CCR^16^) or iteratively testing for regions within genes containing less observed missense variation than expected (MCR^17^). Several approaches have attempted to model site-specific intolerance through a sliding window approach (MTR^18^), homologous residues across protein domains (HMC^19^), or by integrating the 3D structural context of the amino acids (COSMIS^20^).

A limitation of sub-genic approaches to estimating intolerance is that the precision of such estimates is limited by the amount of standing variation present in the population. Thus, smaller regions of genes tend to have highly unstable intolerance estimates. To address this, LIMBR^15^ fits a Bayesian hierarchical model that allows the borrowing of information across regions of the genome to help stabilize estimates. While LIMBR has been shown to add utility in the interpretation of pathogenic missense variation, the approach is inferentially limited, as it relies on simplifying Gaussian assumptions that do not fully match the distributional properties of the data.

In this study, we develop a new framework that enables probabilistic inference of regional intolerance, which we call PRIME, that explicitly reflects the data being modeled. Across all coding regions of the genome, we jointly estimate the probability of a variant being missense in the population via a Bayesian hierarchical model. Like LIMBR, this allows for the borrowing of strength across regions, stabilizing estimates in low sample size regions and greatly reducing bias due to region length. In addition to those advantages, PRIME outputs directly interpretable estimates of regional intolerance, providing a statistically robust framework to test biological hypotheses about how purifying selection acts upon the genome. We demonstrate the added utility of this approach across three applications: (i) prioritizing of novel Mendelian disease-associated genes, (ii) revealing how protein architecture and functional specialization shape regional intolerance in gene families, and (iii) prioritizing variants for pathogenicity in genetic diagnosis.

## Results

### PRIME: Probabilistic Regional Intolerance to Missense Estimation

PRIME is a probabilistic graphical model that utilizes standing variation in the human population to jointly estimate intolerance across all sub-genic regions in the genome. It models variant counts in regions as a binomial process, estimating the probability of a variant in a region being missense. This places the resulting intolerance estimates on a common, interpretable scale, enabling direct comparisons across regions. As a Bayesian model, PRIME captures uncertainty in sub-genic estimates through a joint posterior distribution, allowing this uncertainty to be propagated through all downstream analyses. Furthermore, PRIME models dependencies hierarchically, allowing information sharing across regions and stabilizing estimates in data-sparse settings.

The PRIME model is fit to subregions of protein-coding genes defined by the Consensus Coding Sequence (CCDS) project^21^. For each gene, we selected the Ensembl^22^ canonical transcript to provide a single, standardized representation of exon structure and protein sequence (Methods: Gene Model). Sub-genic regions were then defined under two alternative schemes: protein domain-based regions, corresponding to annotated domain boundaries mapped onto the canonical protein sequence (Fig. 1a), and exon-based regions, corresponding to individual protein-coding exons of the canonical transcript (Supplementary Fig. 1a). Variants from the gnomAD v4.1^4^ exome and genome sequence data were functionally annotated and then counted within each region (Methods: Variant Annotation), allowing PRIME to leverage standing variation observed across 730,947 exomes and 76,215 genomes for inference. on protein domain boundaries, and gnomAD exome and genome variants are counted in each region after consequence annotation. **b**, Graphical model of PRIME, fit to obtained variant counts from previous step. Colored nodes denote observed data and unfilled nodes denote latent parameters.

**Figure 1:**
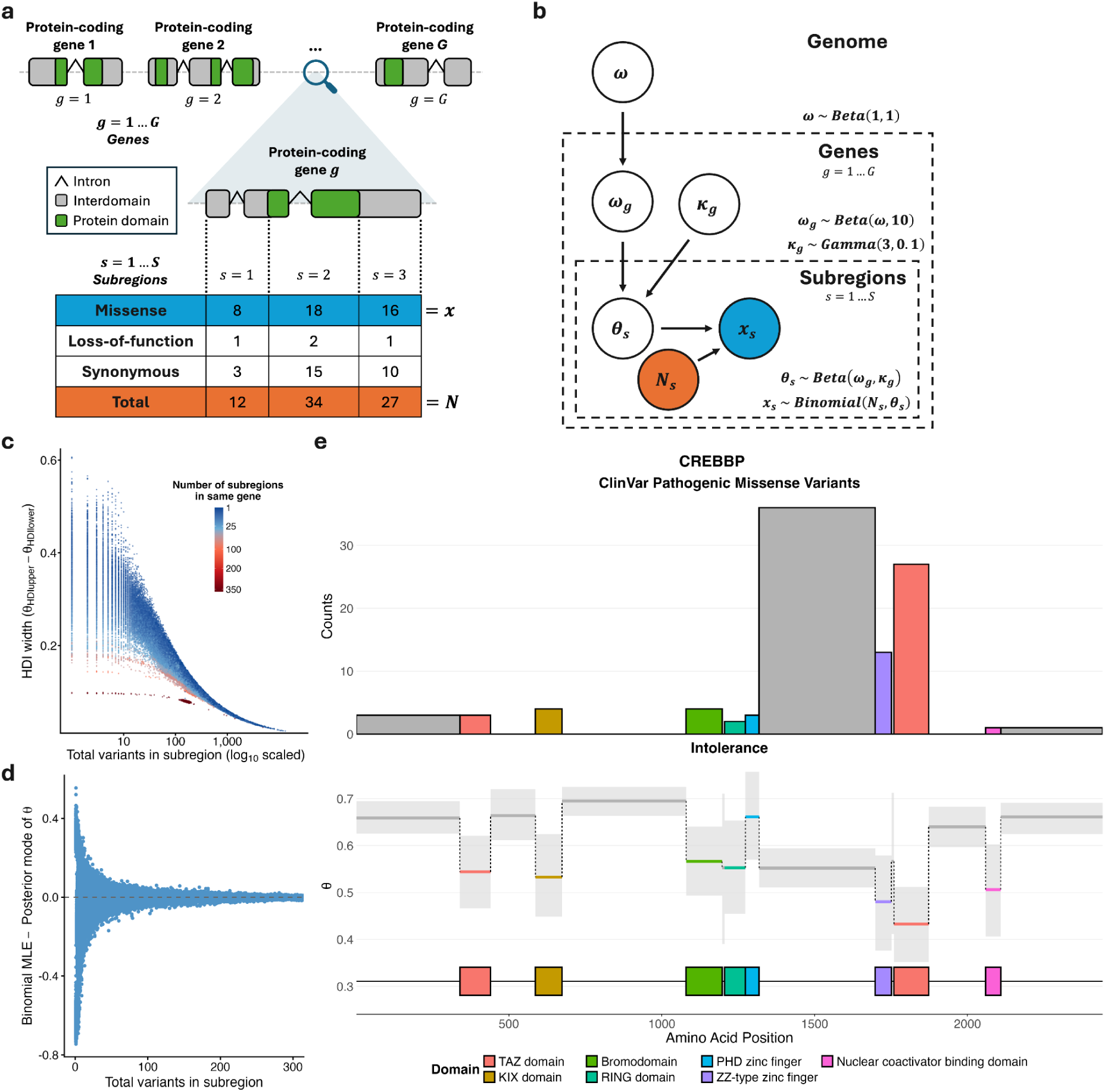
PRIME Framework and Output For Gene with Localized Intolerance. **a**, Workflow to obtain regional variant counts. Protein-coding genes are divided into subregions based

PRIME jointly estimates θ, the probability of a variant in the population within a region being missense. **c**, Uncertainty of θ captured in the marginal posterior distributions. Regions with more standing variation in the population have a tighter distribution that is shown through a smaller 95% highest density interval (HDI) width. Regions with fewer variants have a more diffuse distribution, reflecting greater uncertainty in their estimate of θ. Regions nested in genes with fewer subregions have more uncertainty in their estimates due to a weaker gene-level prior. **d**, Shrinkage effect of hierarchical model on regional estimates of intolerance. Regions with a high number of variants are primarily informed by the variant information within them, whereas lower sample size regions have more influence from the prior leading to a stronger shrinkage effect away from the maximum likelihood estimate. This effect stabilizes estimates when the amount of standing variation in a region is low. **e**, Regional intolerance of disease gene CREBBP captures distribution of pathogenic missense variants. Top: ClinVar pathogenic missense variant counts in each region. Bottom: PRIME regional estimates of the probability of a population variant being missense (θ). Horizontal segments denote the posterior mode of θ for a given region, with gray shading indicating the 95% HDI.

The PRIME model was fit jointly across all subregions within each gene, allowing regional intolerance estimates to be informed by both gene-level and genome-wide intolerance (Fig. 1b, Supplementary Fig. 1b; Methods: Estimating Regional Intolerance with PRIME).

Standing variation was represented as count data and modeled using a binomial likelihood (Equation 1), which naturally reflects the underlying variant-generating process in the population. Within each region, observed missense variants were treated as ‘successes’ and total observed variants as ‘trials’. PRIME thus estimates a region-specific ‘success’ probability, θ, interpreted as the probability that a variant from the population within that region is missense. This parameter is defined conditional on the total number of observed variants in the region, thereby accounting for regional differences in underlying mutability. Lower values of θ, reflecting a depletion of functional variants relative to total in the population, indicate greater intolerance to variation due to stronger purifying selection acting on the region.

The Bayesian hierarchical framework of this approach offers several advantages^15^. Rather than producing a point estimate of θ, PRIME yields posterior distributions that quantify uncertainty in regional intolerance. These posteriors are more concentrated in regions with greater standing population variation and more diffuse in regions with fewer observed variants, reflecting differences in statistical information (Fig. 1c). This is especially important for sub-genic approaches where we are underpowered to measure intolerance in smaller regions due to limited sample size.

As a hierarchical model, PRIME utilizes the natural grouping of regions within genes to pool information across regions. Regional intolerance parameters are modeled through a shared gene-level prior (Equation 2), whose posterior is jointly informed by all regions within the gene. This hierarchical structure induces shrinkage, pulling estimates of θ for regions with limited data toward the gene-level intolerance estimate, which itself is regularized toward a genome-wide prior (Equation 3). As the number of observed variants in a region increases, this influence diminishes, and θ converges to the binomial maximum likelihood estimate – the empirical proportion of missense to total variants within the region itself (Fig. 1d). This hierarchical shrinkage effect stabilizes estimates in regions with limited data, which are often shorter and would otherwise yield noisy estimates.

PRIME infers all parameters jointly, which can each be summarized by their marginal posterior mode and 95% highest density intervals (Supplementary Data 1). These estimates of regional intolerance appear to recapitulate the distribution of pathogenic missense variants in disease-associated genes, which often cluster in intolerant regions^14,15^. We formally demonstrate this relationship across multiple disease genes below, and present the highly modular transcriptional coactivator *CREBBP* as an illustrative example among 347 genes showing a significant correlation between pathogenic variant density and domain-level intolerance (Fig. 1e).

### Pathogenic Missense Variants Are Enriched in Intolerant Regions

Intolerant regions are depleted for functional variation relative to expectation, consistent with stronger purifying selection acting on variants in these regions. Accordingly, deleterious variants are enriched in these regions, as shown from prior genic and sub-genic intolerance metrics^1–4,14–20^. PRIME intolerance estimates similarly prioritize regions harboring ClinVar^23^ pathogenic missense variants in both domain and exon-based delineations of genes across the exome. Domain-based PRIME estimates achieve the highest cumulative enrichment, as measured by area under the curve (AUC), exceeding both subRVIS^14^ scores recomputed in the same regions with the same variant counts and MCRs^17^ across the same set of genes (Fig. 2a). Focusing on the most intolerant 10% of coding sequence for each method, which represent the highest-confidence regions for variant prioritization, PRIME domains capture the largest fraction of pathogenic missense variants (32.4%), compared with PRIME exons (30.7%), MCRs (31.3%), domain-based subRVIS (18.3%), and exon-based subRVIS (13.5%). This enrichment is highly significant relative to the most tolerant regions of coding sequence in PRIME, even after normalizing for length. Domain-based regions show a 7.67-fold enrichment in pathogenic missense variant density between the most and least intolerant deciles (6204.1 vs. 809.0 variants per megabase, exact test p < 2.2e-16), while exon-based regions show a corresponding 6.02-fold enrichment (5562.9 vs. 923.6 variants per megabase, exact test p < 2.2e-16; Supplementary Fig. 2).

**Figure 2:**
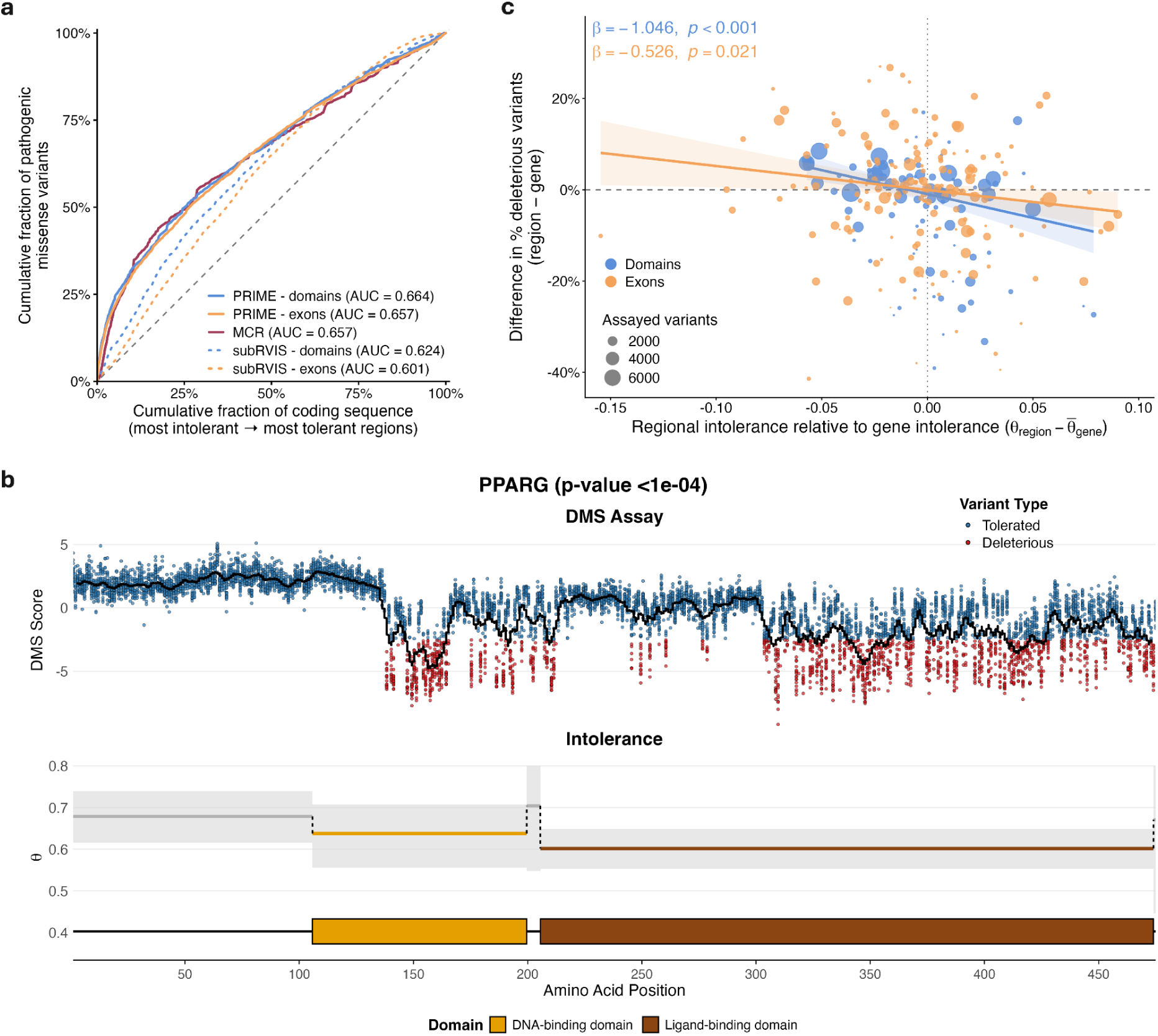
Intolerant Regions Harbor Excess Deleterious Missense Variation. **a**, Cumulative enrichment of ClinVar pathogenic missense variants across regions ranked by intolerance. Coding regions were ordered from most to least intolerant based on PRIME estimates (posterior mode of θ), subRVIS scores, or MCR observed/expected, and the cumulative fraction of coding sequence (x-axis) was plotted against the cumulative fraction of ClinVar pathogenic missense variants (y-axis). Upward direction of curve from null expectation (uniform distribution of variants) indicates enrichment of pathogenic variants in more intolerant regions. AUC for each method is reported in the legend, summarizing how effectively each approach captures pathogenic missense variants across the full ranking of coding sequence. **b**, Regional intolerance of disease gene PPARG correlates with deep mutational scanning (DMS) scores. Top: experimental DMS scores for missense variants across the protein sequence, with variants classified as functionally tolerated (blue) or deleterious (red). The black curve indicates a sliding-window mean. Bottom: PRIME regional estimates with 95% credible intervals of the probability of a variant being missense (θ). **c**, Within-gene enrichment of deleterious DMS variants across regions ranked by intolerance. Regions were ordered from most to least intolerant based on PRIME estimates (posterior mode of θ), and the cumulative fraction of assayed variants (x-axis) was plotted against the cumulative fraction of within-gene excess deleterious variants (y-axis). Excess deleterious variants were defined relative to the gene-specific baseline rate of deleterious variants, controlling for differences in the rate of assayed deleterious variants across genes. Deviation above null expectation of 0 in more intolerant regions indicates enrichment of deleterious variants relative to a gene’s expectation.

To evaluate whether PRIME regional intolerance predicts the localization of pathogenic variants beyond mutational opportunity within genes, we implemented a gene-by-gene ClinVar permutation test in which pathogenic missense variants were redistributed across regions according to cumulative mutation rates (Methods: Association of Regional Intolerance and ClinVar Missense Pathogenicity, Equation 4). 347/2,834 (12.2%) genes for the domain model and 293/3,206 (9.1%) genes for the exon model were significant at ɑ = 0.05 with a Benjamini-Hochberg adjustment, indicating for these genes that pathogenic missense variants are enriched in intolerant regions beyond what would be expected from mutational burden alone (Supplementary Data 2). The transcriptional coactivator gene *CREBBP*, presented in Fig. 1e, is one such example: pathogenic missense variants associated with Menke-Hennekam syndrome 1 (MIM #618332) or Rubinstein-Taybi syndrome 1 (MIM #180849) are concentrated in regions with strong inferred intolerance, such as the ZZ-type zinc finger and TAZ2 domains. Interestingly, the disruption of these two critical domains has even been associated with distinct disease subtypes^24^.

While the above permutation approach controls for regional mutation rate differences, it can not account for unknown ascertainment biases in ClinVar, where pathogenic variants may be preferentially identified and reported in functionally characterized regions of genes. Thus, we extended this permutation framework to assess whether our regional intolerance estimates are predictive of experimentally measured functional impact using deep mutational scanning (DMS) data from ProteinGym^25^ (Methods: Association of Regional Intolerance and Experimental DMS Assay Scores, Equation 5). 10/27 (37.0%) genes demonstrated a significant association between PRIME domain regional intolerance and experimentally measured deleterious effects in deep mutational scanning assays, where 11/30 (36.7%) demonstrated significance for exon regional intolerance at ɑ = 0.05 with a Benjamini-Hochberg adjustment (Supplementary Data 3). One such gene is the nuclear receptor *PPARG*, in which more deleterious DMS scores are observed in the highly conserved^26^ DNA-binding and ligand-binding domains, aligning with the inferred pattern of regional intolerance (Fig. 2b). To assess whether intolerant regions within genes are consistently enriched for deleterious variants across ProteinGym assays, we estimated each gene’s baseline deleterious-variant rate and quantified the deviation from this baseline for each region (Methods: Association of Regional Intolerance and Experimental DMS Assay Scores). Regions that were more intolerant than their gene-wide mean had higher-than-expected deleterious-variant rates, whereas relatively tolerant regions had lower-than-expected rates, with significant negative associations at ɑ = 0.05 for both domain and exon-based regions (Fig. 2c).

Using the results from the ClinVar permutation test, we tested for enrichment of several Online Mendelian Inheritance in Man (OMIM)^27^ disease gene sets (Supplementary Data 4), representing different modes of disease inheritance, to assess whether specific inheritance mechanisms are overrepresented among the set of significant genes (Methods: Enrichment Testing of Gene Sets in ClinVar Permutation Test). All OMIM gene sets except autosomal recessive disease-linked genes showed significant enrichment in both domain and exon-based versions of the test (Table 1). The failure to show enrichment in recessive disease genes was expected, as variants causing autosomal recessive disorders are typically tolerated in heterozygous carriers and thus are subject to weaker selection^28^. We then followed a similar approach for all Hugo Gene Nomenclature Committee (HGNC)^29^ gene groups, extending the analysis to a more comprehensive set of genes grouped by shared function and evolutionary relationships rather than disease annotation alone. Given the hierarchical nature of these groupings, we implemented a gatekeeping procedure^30,31^ to control Type I error from the multiple enrichment tests, testing a gene group only if their parent group was significantly enriched (Supplementary Data 5; Methods: Enrichment Testing of Gene Sets in ClinVar Permutation Test). 24 groups showed significance after multiple test corrections with the domain-based regions (Fig. 3a), and 27 did for exon-based regions (Fig. 3b). Many larger gene groups such as ion channels, chromatin modifying enzymes, and zinc fingers showed significant enrichment from both versions of the tests. Interestingly, some gene groups showed significance exclusively in only one delineation of genes. The homeobox transcription factors were only significant for domains (Holm-adjusted p = 3.26e-03), whereas GABA receptors and ATP-dependent chromatin remodeling complexes were only significant for exons (Holm-adjusted p = 5.90e-04 and p = 8.10e-03, respectively).

**Figure 3:**
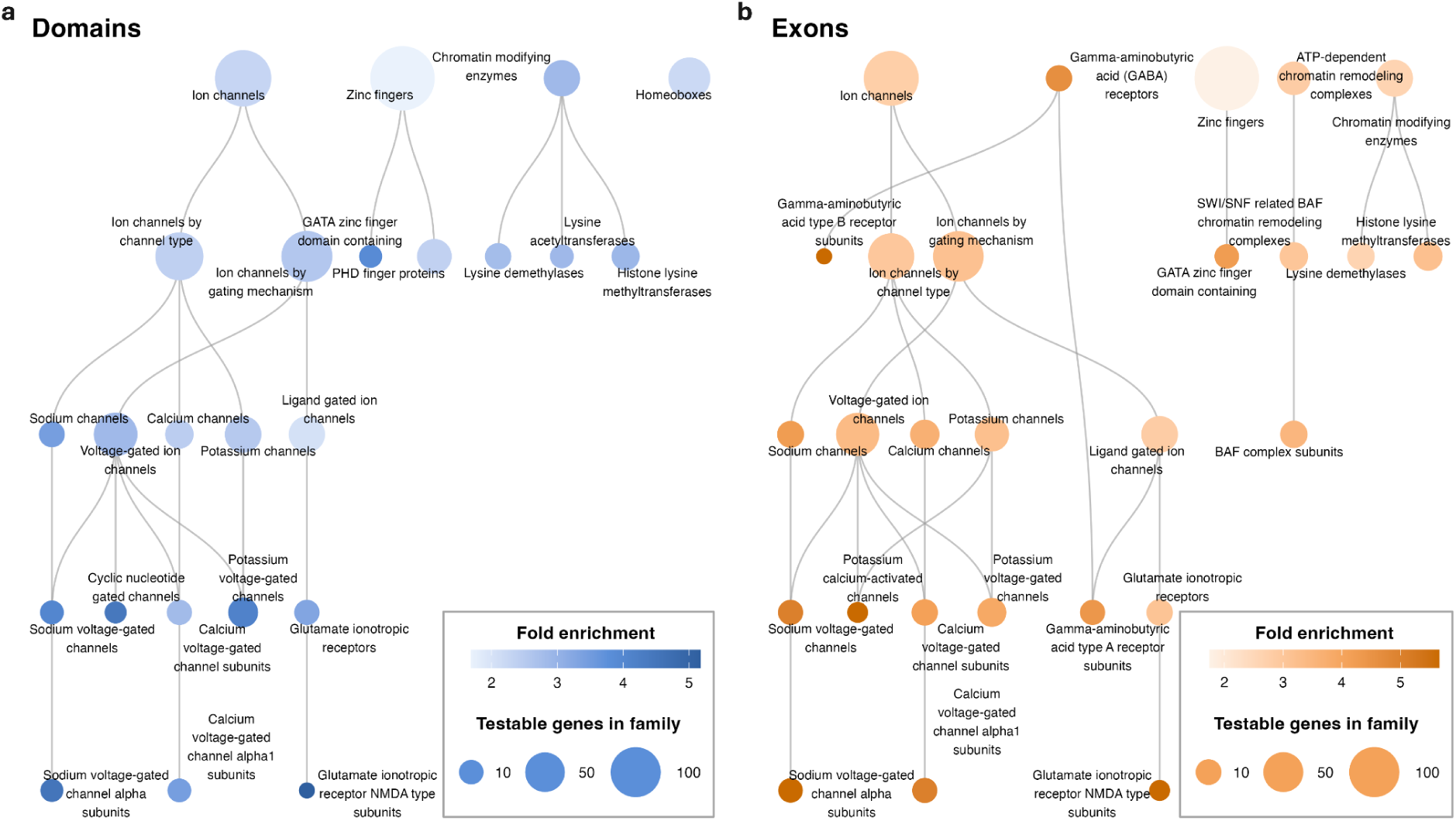
HGNC Gene Group Enrichment in ClinVar Permutation Test. **a**, Tree of HGNC gene groups that show significant enrichment in domain-based gene-level permutation tests. **b**, Tree of HGNC gene groups that show significant enrichment in exon-based gene-level permutation tests. Significance for both was determined by a hierarchical gatekeeping procedure that controls Type I error from multiple testing.

**Table 1:** Enrichment of OMIM Gene Sets in ClinVar Permutation Test. Different OMIM-defined Mendelian disease gene sets and their enrichment for genes with significant association between their domain-based and exon-based regional intolerance and the distribution of their ClinVar pathogenic variants. P-values were computed using an exact test, and asterisks indicate statistically significant enrichment.

| OMIM Gene Set | Domains |  |  |  | Exons |  |  |  |
| --- | --- | --- | --- | --- | --- | --- | --- | --- |
|  | Number of Testable Genes | Number of Significant Genes | Fold Enrichment | Exact p-value | Number of Testable Genes | Number of Significant Genes | Fold Enrichment | Exact p-value |
| De Novo | 971 | 348 | 1.85 | 2.18e-55* | 1049 | 355 | 1.91 | 3.94e-59* |
| Autosomal Dominant | 1124 | 364 | 1.67 | 2.18e-45* | 1200 | 373 | 1.76 | 8.30e-52* |
| Dominant-Negative | 513 | 181 | 1.82 | 1.23e-21* | 536 | 188 | 1.98 | 4.15e-27* |
| Gain-of-Function | 379 | 146 | 1.99 | 3.16e-21* | 392 | 149 | 2.15 | 5.22e-25* |
| Haploinsufficient | 192 | 85 | 2.29 | 1.97e-16* | 218 | 78 | 2.02 | 2.47e-11* |
| All OMIM | 2509 | 523 | 1.08 | 5.65e-10* | 2847 | 543 | 1.08 | 1.51e-10* |
| Autosomal Recessive | 1582 | 205 | 0.67 | 1.00e+00 | 1846 | 213 | 0.65 | 1.00e+00 |

### The Weakest Link: Characterizing Genes by Their Most Intolerant Region

Analyses of population variation have shown that some genes are markedly depleted for functional variants, a signal captured by genic intolerance metrics and commonly used to prioritize candidate disease genes^1–4^. However, gene-level intolerance scores can underestimate the functional importance of genes with pronounced regional heterogeneity in intolerance, as tolerant regions will tend to dilute or mask highly intolerant regions. An example is *FOXC1* (Fig. 4a), which has a negative gene-level missense Z-score^2^, suggesting the gene is tolerant to missense variation. However, *FOXC1* is an established disease gene with a concentration of known pathogenic missense variants lying within its DNA-binding forkhead domain^32–34^. PRIME captures the strong intolerance of this domain and, given the tolerance of the surrounding coding sequence, an explanation for why missense Z categorized the gene as tolerant: the majority of the gene was tolerant and only the relatively small forkhead domain showed a substantial depletion of missense mutation. This example illustrates how regional heterogeneity can dilute gene-level intolerance metrics leading to misclassification. It also illustrates an important principle: disease relevance is not due to an average property of a gene but of the extremes. If only a small region of the gene harbors pathogenic variants, the gene is disease relevant, even if the majority of the gene is flooded with benign variation. We leverage this principle to systematically characterize genes’ disease relevance by focusing on their ‘weakest link’ - the region most sensitive to structural disruption and therefore most likely to lead to disease when mutated.

**Figure 4:**
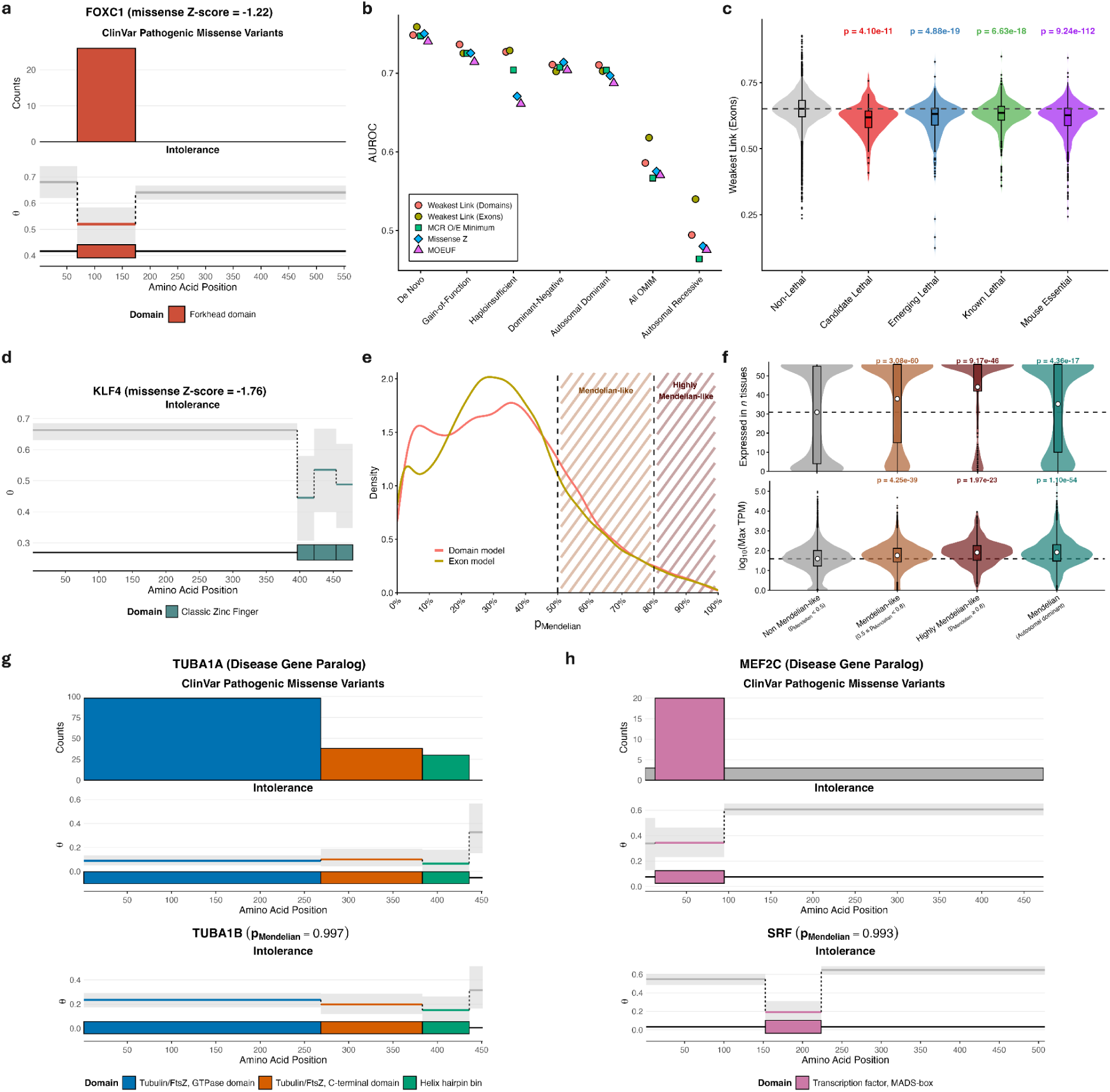
The Weakest Link Statistic Discriminates Disease Genes and Prioritizes Disease-Causing Gene Candidates. **a**, Regional intolerance of disease gene *FOXC1* captures localized intolerance in the forkhead domain. Gene ranks highly among all genes based on weakest link intolerance (95th percentile), despite having a negative missense Z-score indicating tolerance to missense variants with more observed in the population across the entire gene than expected. ClinVar pathogenic missense variants are highly concentrated in the intolerant forkhead domain. **b**, AUROC comparison of the weakest link to missense constraint metrics for discriminating sets of established disease genes in the OMIM database. Both domain (red circle) and exon (yellow circle) derived weakest link statistics are consistently amongst the top performing methods. **c**, Distribution of the exon-based weakest link statistic for lethal genes with varying levels of evidence and genes essential for development in mice. Above p-values are from one-sided Mann-Whitney tests comparing the distributional differences between the lethal/essential gene sets and the complement of those sets combined (non-lethal genes). Dashed line represents the median weakest link value among non-lethal group. **d**, Regional intolerance of non-disease gene *KLF4* captures localized intolerance in the three zinc finger C-terminal DNA-binding domain. Gene ranks highly among all non-disease genes based on weakest link intolerance (99th percentile) with a negative missense Z-score. **e**, Density of both domain and exon-based p_Mendelian_ statistic across all non-OMIM genes. Genes with values above 0.5 are Mendelian-like, displaying localized intolerance stronger than that of most known autosomal dominant Mendelian disease-associated genes. Genes with values above 0.8 are highly Mendelian-like, with stronger localized intolerance than the vast majority of autosomal dominant Mendelian genes. **f**, Mendelian-like genes show more ubiquitous expression across tissues and higher expression in specific tissues compared to non-Mendelian genes, consistent with expression patterns in known disease genes. Top: Number of tissues a gene is expressed in (median TPM >3 threshold used). Bottom: Log_10_ scaled maximum expression across tissues, defined as the maximum of median TPM values per tissue. Dashed lines represent the mean value in the non Mendelian-like set. **g**, Regional intolerance of *TUBA1B* (gene with the highest p_Mendelian_) and disease gene paralog *TUBA1A*. Genes display a similar pattern of localized intolerance that captures distribution of pathogenic ClinVar variants in *TUBA1A*. **h**, Regional intolerance of *SRF* (gene with the second highest p_Mendelian_) and disease gene paralog *MEF2C*. Intolerance is concentrated in the MADS-box domain of both genes, where pathogenic missense variants cluster in *MEF2C*.

For each gene, we define the weakest link as the minimum intolerance value across all of its subregions. Usually, such a parameter would be difficult to estimate, as the minimum would be dominated by small, highly unstable subregions. This illustrates the practical utility of our Bayesian model which elicits a joint posterior over all subregions. To estimate the weakest link, we simply take the minimum across each gene for each Markov Chain Monte Carlo (MCMC) iteration (Methods: The Weakest Link of Genes). Each MCMC iteration is a sample from the joint posterior distribution, and by computing the minimum across samples we get the posterior of the weakest link statistic for each gene. This procedure characterizes the full posterior of each genes’ most intolerant subregion while explicitly propagating subregional uncertainty into the distribution. We summarize each gene-level posterior distribution of minimums by taking the upper bound of its 95% highest density interval. Since higher values are less intolerant, this incorporates the uncertainty into the summary, giving a conservative summary statistic of the gene’s weakest link. This statistic was computed separately using both domain and exon-based partitions of genes (Supplementary Data 6). We demonstrate its utility as a gene-level intolerance measure by evaluating its ability to discriminate several OMIM disease gene sets and lethal/essential genes. Relative to previously established regional^17^ and gene-level^2,4^ missense constraint metrics, the weakest link provides superior discrimination of disease genes across most evaluated OMIM gene sets (Fig. 4b, Supplementary Table 1). We additionally compared the weakest link to RVIS and the gene-level intolerance parameter from PRIME (ω*_g_*), both estimated from the same underlying variant counts, demonstrating that prioritizing the most intolerant region provides greater discriminatory power than gene-level aggregation (Supplementary Table 1). Further, lethal genes^35^ and mouse essential genes^36,37^ are significantly more intolerant than non-lethal genes based on their weakest link statistic (Fig. 4c, Supplementary Fig. 3).

Genes associated with Mendelian disorders exhibit an especially strong depletion of functional variants in their most intolerant subregions, or weakest links, particularly for autosomal dominant disorders where a single heterozygous missense variant in a functionally critical region can be pathogenic. Interestingly, we also see non-Mendelian genes that, on a gene level, are tolerant, but contain subregions that are highly intolerant (see Fig. 4d for example). We therefore asked how common these non-Mendelian genes with highly intolerant subregions comparable to Mendelian genes were across the genome. To quantify this, we constructed a reference distribution of weakest link intolerance across all known autosomal dominant Mendelian genes and compared each non-Mendelian gene to this distribution. For each gene, we computed p_Mendelian_ (Equation 6), the posterior mean probability that its weakest link is more intolerant than that of a randomly sampled autosomal dominant Mendelian gene. This statistic was computed using both domain and exon-based partitions of genes (Supplementary Data 6), providing a quantitative measure of how closely a gene’s localized intolerance resembles that of known disease genes.

Surprisingly, we find 462/17,143 genes (2.7%) that are currently not associated with an autosomal dominant disease phenotype in the OMIM database have a domain-based p_Mendelian_ above 0.8, and 3,688/17,143 (21.5%) above 0.5 (Fig. 4e). This indicates that these genes display even stronger selection on their weakest link than most known dominant disease genes, providing evidence towards potential undiscovered disease mechanisms from their disruption. Using mouse essential genes as an orthogonal control, many of which lack annotated Mendelian phenotypes in humans, we found a significant enrichment of these genes among those with high p_Mendelian_ values. Essential genes were 1.98-fold enriched among non-Mendelian genes with p_Mendelian_ ≥ 0.8 compared to all non-Mendelian genes (exact test p = 4.73e-09), suggesting that this metric is capturing biologically important intolerance beyond currently annotated disease genes.

Another way of assessing the biological importance of genes is through the breadth of their expression across tissues and the magnitude of their expression within specific tissue types. Previous work has shown that genes constrained for missense and loss-of-function variation exhibit significantly higher and broader expression than less constrained genes when using gene-level metrics^3,4^. To assess whether weakest link intolerance captures the same relationship, we similarly used GTEx^38^ median gene-level TPM data to compare expression across groups of genes stratified by weakest link intolerance (Methods: The Weakest Link of Genes). Mendelian-like genes (0.5 ≤ p_Mendelian_ < 0.8) and highly Mendelian-like genes (p_Mendelian_ ≥ 0.8) show significantly broader expression across tissues and higher expression in specific tissue types compared to genes with weak evidence of being Mendelian-like (p_Mendelian_ < 0.5), consistent with observed expression patterns in known autosomal dominant disease genes (Fig. 4f, Supplementary Figs. 4,5).

Interestingly, we find significant overlap in the function of genes with the top domain-based p_Mendelian_ values, with 7 of the top 20 being related to RNA processing, and 4 of the top 20 involved in proteostasis. This enrichment is consistent with the modular architecture of these systems, where essential functional elements - such as RNA-binding motifs or catalytic interfaces - can be highly intolerant despite greater tolerance to variation elsewhere in the gene. Further, 9 of the top 20 genes also have at least one paralogous gene that is currently implicated in autosomal dominant disease in OMIM (Table 2).

**Table 2:**
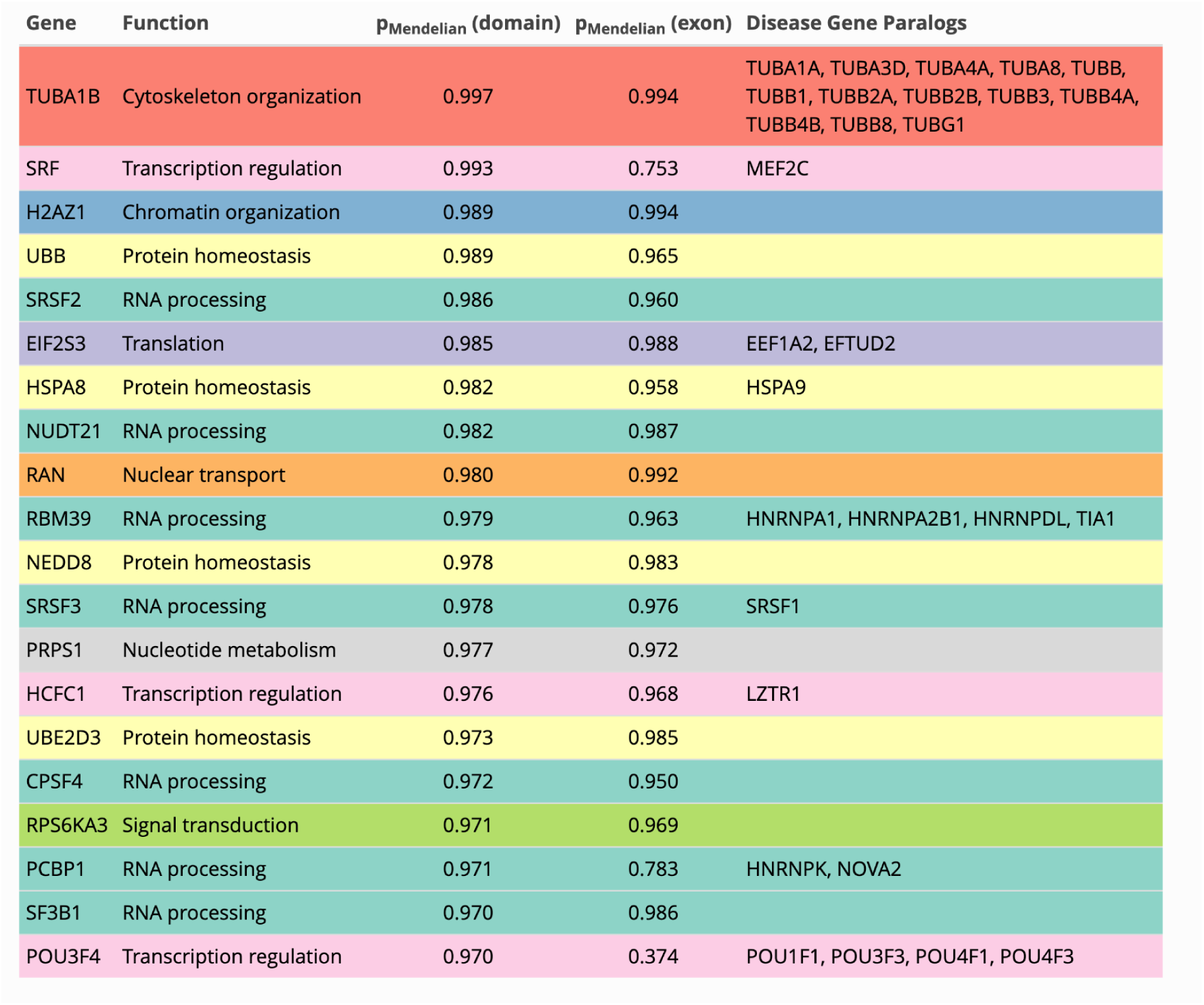
Top 20 Mendelian-like Genes. Set of top 20 genes that display strongest signals of domain-based localized intolerance compared to known autosomal dominant Mendelian disease genes. Gene paralogs that have an autosomal dominant disease phenotype in OMIM are listed in the final column. Gene functions were assigned by manual curation.

The gene with the highest p_Mendelian_ domain value (0.997) among all non-OMIM genes is the alpha-tubulin gene *TUBA1B*, which is involved in microtubule cytoskeleton organization. This gene is expressed across all 56 tissue types considered in the GTEx dataset and also has a maximum tissue-specific TPM of 1041.74 (98th percentile of non-OMIM genes), suggesting both broad tissue expression and high peak expression. Despite its strong regional intolerance and expression in tissues, *TUBA1B* has not been associated with a Mendelian phenotype.

While *TUBA1B* in its entirety is intolerant to missense variation, its helix hairpin bin (amino acids 384-436) is particularly depleted of missense variation, with only 2 missense variants out of 30 total variants observed in gnomAD. A paralog of *TUBA1B*, *TUBA1A*, shares 99.5% sequence identity with *TUBA1B* and differs by only two amino acids^39^. *TUBA1A* encodes the majority of α-tubulin in developing brains and is known to harbor pathogenic mutations that cause lissencephaly (MIM #611603). Both genes display similar patterns of the strongest intolerance occurring in protein domains, particularly in the helix hairpin bin (Fig. 4g). In *TUBA1A*, 30 of 166 (∼18%) ClinVar pathogenic missense variants fall within the helix hairpin, despite this region constituting only ∼12% of the amino acids of the encoded protein.

The serum response factor gene *SRF*, which has the second highest p_Mendelian_ domain value (0.993), encodes a ubiquitous MADS-box transcription factor that plays a key role in transcriptional programs controlling cytoskeletal organization and muscle development. Deletion of *SRF* is lethal in a variety of organisms across multiple phyla, including mice, in which loss of *SRF* causes incomplete gastrulation and embryonic arrest^40^. *SRF*, like *TUBA1B*, is also expressed in the maximum possible number of tissue types, and has a maximum tissue-specific TPM of 165.05 (84th percentile of non-OMIM genes), supporting its role as a ubiquitous transcription factor. Intolerance is strongest in the DNA-binding MADS-box domain (amino acids 153-223) of this gene, which is highly depleted of missense variants relative to the rest of the gene (Fig. 4h). The same pattern can be seen in the disease gene paralog of *SRF*, *MEF2C*, where pathogenic variants are associated with 5q14.3 microdeletion syndrome and neurodevelopmental disorder with hypotonia, stereotypic hand movements, and impaired language (MIM #613443). Additionally, 20 of 26 (∼77%) of ClinVar pathogenic missense variants fall in the MADS-box domain of *MEF2C*, which is only ∼17% of the total protein sequence.

### Functional Domains Shape Patterns of Purifying Selection Within Gene Families

Gene families arise through duplication and divergence^41^, often retaining conserved domain architectures that underlie shared molecular functions^42,43^. Because these domains encode core structural and biochemical properties, they are frequently preserved across paralogs and thus may be under a similar degree of selection across these genes. Consistent with this, several HGNC gene families sharing common domain architectures were enriched for genes in which the regional distribution of intolerance was significantly associated with the distribution of pathogenic missense variants (Fig. 3a). Investigating such patterns of selection in related genes may yield insights into the evolutionary preservation of critical molecular functions. The probabilistic framework and joint posterior distribution across genes of PRIME allows us to formally interrogate such relationships. With regional intolerance estimates placed on a consistent, interpretable probabilistic scale across all genes, we can perform direct quantitative comparisons within and across gene families. Critically, these comparisons can leverage the entire posterior distribution, incorporating uncertainty in the analyses rather than relying on point estimates.

We first demonstrate the application of this framework in voltage-gated potassium (Kv) channels, the largest subset of potassium channel genes, comprising 40 genes in total. Kv channels respond to changes in membrane potential through voltage-dependent gating and are critical regulators of electrical signaling in excitable cells, particularly neurons. Mutations in these genes have been implicated in a wide variety of neurological disorders, including epilepsy, ataxia, autism spectrum disorder, and schizophrenia^44^. Further, voltage-gated potassium channels were significant in the HGNC enrichment analysis (Fig. 3a, Holm-adjusted p = 3.54e-08), demonstrating that regional intolerance captures the distribution of these pathogenic variants across the gene family. The structure of these proteins is highly conserved, with all members containing six transmembrane ɑ-helical segments S1-S6, where the first four S1-S4 segments act as the voltage-sensing domains and the last two S5-S6 segments conduct potassium ions^45^. Despite being connected through these segments, the function of these two domains is highly independent^46^ - the voltage-sensing domain can even be expressed as an isolated domain^47^ or transferred to a non-voltage-gated channel to allow voltage sensing^48,49^.

The highly modular organization of these key domains suggests that they may be under different selective constraints. To formally interrogate this across the gene family, we first identified 28 Kv genes with consistent protein domain annotations corresponding to these two distinct domains. Using PRIME, we then computed, at each MCMC iteration, the difference in the probability of a variant being missense between the pore-forming and voltage-sensing domains within each Kv gene, thereby generating a posterior distribution of this within-gene difference (Fig. 5a). All genes but one have a negative posterior mean difference, meaning their pore-forming domain is more intolerant to variation than their voltage-sensing domain, with 10 demonstrating a negative difference in at least 97.5% of their MCMC samples (Supplementary Data 7). Collectively, these results indicate that the ion-conducting pore region is under stronger purifying selection than the voltage-sensing domain across this gene family. Notably, the magnitude of this difference varies according to subunit function. Some Kv genes encode modulatory subunits that are not electrically functional on their own but instead coassemble with conducting subunits to form heterotetrameric channels, where they modulate channel current^50^. We observed a significantly weaker effect among these silent subunits (Fig. 5a), with a posterior probability of 0.9963 that the mean (θ_pore-forming_ - θ_voltage-sensing_) difference was greater in magnitude for conducting than for silent subunits (Supplementary Fig. 6a).

**Figure 5:**
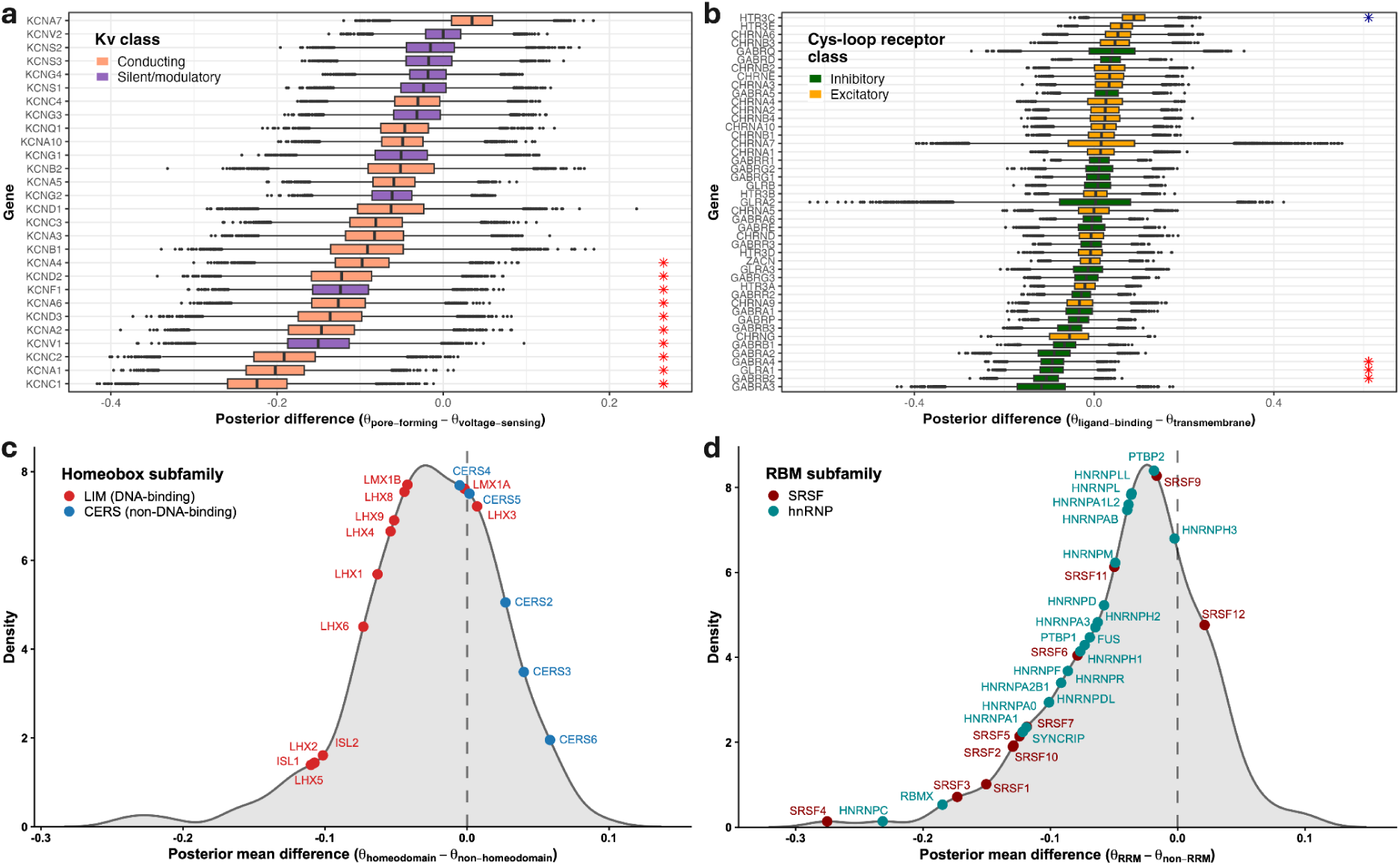
Gene Families Demonstrate Consistent Patterns of Purifying Selection in Shared Functional Domains. **a**, Voltage-gated potassium channels show stronger intolerance in the pore-forming domain than the voltage-sensing domain. The difference in θ was computed between the two regions at each MCMC sample for every Kv gene, with a negative difference indicating stronger intolerance in the pore-forming domain. Red asterisks indicate genes for which at least 97.5% of posterior samples of the within-gene difference were below 0 (stronger pore-forming intolerance), whereas blue asterisks indicate genes for which at least 97.5% of samples were above 0 (stronger voltage-sensing intolerance). Conducting Kv channels show a stronger effect compared to silent/modulatory subunits. **b**, Ligand-binding domain intolerance compared to transmembrane domain intolerance in ligand-gated ion channels. Red asterisks indicate genes for which at least 97.5% of posterior samples of the within-gene difference were below 0 (stronger ligand-binding intolerance), whereas blue asterisks indicate genes for which at least 97.5% of samples were above 0 (stronger transmembrane intolerance). Inhibitory subunits show stronger intolerance on average in the ligand-binding domain, where excitatory subunits show stronger intolerance on average in the transmembrane domain. **c**, Homeobox genes demonstrate consistent localized intolerance in the DNA-binding homeodomain. Density of the posterior mean difference in θ between the homeodomain regions and non-homeodomain regions for all homeobox genes is shown. Distribution is shifted below 0, indicating stronger intolerance in the homeodomain in most genes. **d**, RNA-binding motif (RBM) genes demonstrate consistent localized intolerance in the RNA-recognition motif (RRM). Density of the posterior mean difference in θ between the RRM regions and non-RRM regions for all RBM genes is shown. Distribution is shifted below 0, indicating stronger intolerance in the RRM in most genes.

Ligand-gated ion channels also conduct ions, but do so allosterically as a response to agonist binding rather than from changes in membrane potential. Their subunits contain two critical functional domains: a transmembrane domain containing the ion pore and an extracellular domain mediating ligand recognition^51^. These are targets for several drugs and are associated with many central nervous system diseases, such as epilepsy, Alzheimer’s disease, schizophrenia, and Parkinson’s disease^52^. The Cys-loop superfamily of these ion channels in particular are key for proper neuronal activity, mediating both neuronal inhibition and excitation^53^. Both the extracellular ligand-binding and transmembrane pore-forming domains of these subunits have independently been implicated in disease and pharmacological intervention, with pathogenic variants and therapeutic compounds targeting either one^54–58^, motivating an investigation into whether they are under different degrees of selective pressure within these genes. The significant enrichment of the broader ligand-gated ion channel family in the HGNC analysis (Fig. 3a, Holm-adjusted p = 3.05e-03) suggests that the differences in intolerance across these regions are biologically meaningful and correspond to the distribution of pathogenic variation in the genes. We thus followed the same procedure as the previous voltage-gated potassium channel analysis, computing the difference in probabilities across MCMC samples between the ligand-binding extracellular domain and transmembrane pore-forming domain for the 45 Cys-loop receptor genes (Fig. 5b). Interestingly, we find evidence of this effect differing based on the action of the subunits: 16 of 23 (69.6%) inhibitory genes, which allow anion influx into the cell, have a negative posterior mean difference, indicating that their ligand-binding domain is the more intolerant of the two, with 3 of 23 genes exhibiting a negative difference in at least 97.5% of MCMC samples. In contrast, 15 of 22 (68.2%) excitatory genes, which allow cation influx, have a positive posterior mean difference, consistent with the transmembrane domain being more intolerant, with 1 of 22 genes exhibiting a positive difference in at least 97.5% of MCMC samples (Supplementary Data 7). Aggregating across genes by comparing the mean (θ_ligand-binding_ - θ_transmembrane_) between groups, we estimate a posterior probability of 0.9994 that the difference is lower in inhibitory vs. excitatory subunits (Supplementary Fig. 6b). Previous work has suggested that the inhibitory Cys-loop receptors share a similar modality of agonist binding in their extracellular domain due to sequence conservation along with supporting structural and functional evidence^59^, potentially explaining why this domain appears systematically more intolerant in this class compared to in excitatory receptors.

As both Kv channels and Cys-loop receptors are implicated in various neurological disorders, we next tested whether the distribution of pathogenic missense variants in their key functional regions correlates with their inferred intolerance. To do so, we applied the same gene-by-gene ClinVar permutation procedure previously described (Equation 4), but restricted the analysis to the relevant domains within each gene. Specifically, pathogenic missense variants were permuted between the pore-forming and voltage-sensing regions for individual Kv channels, and between the ligand-binding and transmembrane regions for Cys-loop receptors, in proportion to their expected mutational burden. Among Kv genes, 8 of 12 with at least one pathogenic missense variant in either region were significant at α = 0.05, consistent with enrichment of pathogenic variants in the more intolerant pore-forming region. 4 of 19 Cys-loop receptor genes reached significance, indicating a stronger correlation for Kv channels by comparison.

The homeobox gene family is a set of transcription factors characterized by the presence of a highly conserved DNA-binding homeodomain. They play a key role in embryonic development, where mutations can cause severe developmental defects resulting in entire body structures lost or altered^60,61^. As master regulators of developmental processes, we hypothesized that the DNA-binding homeodomain, which mediates the core function of these proteins, is under stronger purifying selection than rest of the gene. The homeoboxes’ significant enrichment in the HGNC analysis (Fig. 3a, Holm-adjusted p = 3.26e-03) further suggests that intolerance may be localized in this key binding domain, where disruption is most likely to impair protein function. Leveraging the full joint posterior distribution of PRIME, we tested this by computing all pairwise differences in the probability of a variant being missense between homeodomain and non-homeodomain regions across the 227 homeobox genes in our dataset. Doing so across all MCMC samples generates a posterior distribution of differences for each gene, which can be summarized by their means (Fig. 5c). Of the 227 homeobox genes, 154 (67.8%) exhibit a posterior mean difference below 0, with 13 genes showing a negative difference in at least 95% of posterior samples, consistent with intolerance concentrated in the homeobox motif (Supplementary Data 7). The magnitude of this effect can vary by class, indicative of differences in functional roles across these proteins. For example, LIM homeobox proteins have highly similar homeodomains relative to those of other homeodomain proteins, suggesting the evolution of a distinct DNA-binding specificity^62^. By comparison, the CERS subfamily is highly divergent from other homeobox genes, encoding transmembrane proteins in which the homeodomain appears non-essential for function and may even be dispensable^63,64^. Computing the mean (θ_homedomain_ - θ_non-homeodomain_) in both subfamilies across MCMC samples, we calculate a posterior probability of 0.9999 that the LIM proteins have a lower mean difference than the CERS proteins (Supplementary Fig. 6c), reflecting the functional divergence of the homeodomains between these classes.

The RNA-binding motif (RBM) gene family is made up of 213 genes encoding RNA-binding proteins, many of which contain RNA recognition motifs (RRMs). These proteins play central roles in RNA metabolism, including splicing, stability, and translation^65^. Their dysfunction has also been implicated in a wide range of human developmental and neurological genetic disorders^66,67^. Given the primary function of most of these proteins requires recognizing and binding RNA, we investigated whether the RRM is under stronger selection than the rest of the protein across this gene family, similarly to the DNA-binding domain of the homeoboxes. To test this, we performed an analysis analogous to that used for the homeobox gene family, computing the posterior mean of all pairwise differences in the probability of a variant being missense between the RRM domain and the non-RRM regions for the 194 RBM genes with an RRM annotation in our dataset (Fig. 5d). 149 of 194 (76.8%) RBM genes have a posterior mean difference below 0, with 11 genes demonstrating a negative difference in at least 95% of MCMC samples, indicating very strong localized intolerance in the RRM (Supplementary Data 7). Notably, the SRSF and hnRNP subfamilies of RBM proteins are prominently represented among these genes, with all but one gene (*SRSF12*) in each subfamily exhibiting a negative posterior mean difference (Fig. 5d). These proteins play central roles in pre-mRNA splicing and are key mediators of splice site recognition: SRSF proteins bind exonic splicing enhancers to promote exon inclusion, whereas hnRNP proteins bind splicing silencers to promote exon skipping^68^. *SRSF12*, which does not exhibit especially strong selection on its RRM, is dispensable in mouse knockout studies when this domain is disrupted by frameshift mutations^69^, consistent with our findings.

### Regional Intolerance Enables Discrimination of Pathogenic and Benign Variants

Missense variants of uncertain significance (VUSs) still pose a major challenge in medical genetics^70^. Interpretation of the functional consequence of missense variants, and in turn, the likelihood for them to cause disease is more ambiguous than for variants predicted to cause loss of function entirely in the encoded protein (e.g. nonsense, frameshift, splice-altering)^71,72^. To aid in the interpretation of these missense variants at scale, many in silico variant pathogenicity predictors have been developed^17,73–86^. To translate their use into a clinical setting, a Bayesian framework^87,88^ has been developed to calibrate these predictors at different thresholds to correspond to different strengths of evidence for pathogenicity or benignity according to the American College of Medical Genetics and Genomics and the Association for Molecular Pathology (ACMG/AMP) guidelines^89^.

Deleterious missense variants in disease genes are rarely distributed uniformly along the protein sequence. Instead, they often cluster within functionally critical regions^5–13^. PRIME can help identify these critical regions through their signatures of purifying selection, providing information that is complementary to most variant-level predictors. A variant may be predicted to strongly disrupt protein function, but such molecular effects do not necessarily translate into reduced organismal fitness or disease in humans. This principle was demonstrated with the first intolerance method, RVIS^1^, where combining RVIS percentiles with PolyPhen-2^73^ scores improved the identification of disease-causing de novo mutations in individuals with severe intellectual disability, epileptic encephalopathies, and autism spectrum disorders. Rule PM1 of the ACMG/AMP guidelines assigns evidence of pathogenicity to variants if they fall in a mutational hotspot and/or critical and well-established functional domain that is lacking benign variation^89^. Regional intolerance estimates offer a quantitative framework for identifying such regions and can therefore support application of the PM1 criterion^90^. This information is orthogonal to rule PP3, which uses evidence from in silico variant predictors, provided these predictors do not use intolerance or population variant data directly in their training.

Given we have a continuous measure of regional intolerance bounded from 0 to 1 with PRIME, we can calibrate our estimates as described by Pejaver et al.^88^ to establish thresholds of pathogenic and benign evidence for missense variants when applying rule PM1. Each variant falls within a coding region that has an estimated probability of a variant being missense and therefore can be assigned the intolerance estimate of that region. Estimates are significantly lower for pathogenic variants than for benign variants in both the ClinVar 2019 calibration set and 2020 validation set provided by Pejaver et al. (Fig. 6a,b), demonstrating that pathogenic variants tend to occur in more intolerant regions in the genome compared to benign variants. To calibrate PRIME, we used the 2019 dataset and computed local posterior probabilities of pathogenicity and benignity at different probability thresholds (Methods: Calibration of PRIME Estimates for Clinical Variant Interpretation). To utilize the uncertainty provided in the posterior distribution for each regional estimate, the upper bound of the 95% highest density interval (θ_HDIupper_) was calibrated for pathogenicity and the lower bound (θ_HDIlower_) for benignity. As lower values of θ indicate more intolerance and higher values more tolerance, they function as conservative estimates of pathogenicity and benignity, respectively. After calibration using 10,000 bootstrap iterations, both bounds reached moderate levels of evidence for their respective variant classes (Supplementary Fig. 7a,b, Supplementary Table 2).

**Figure 6:**
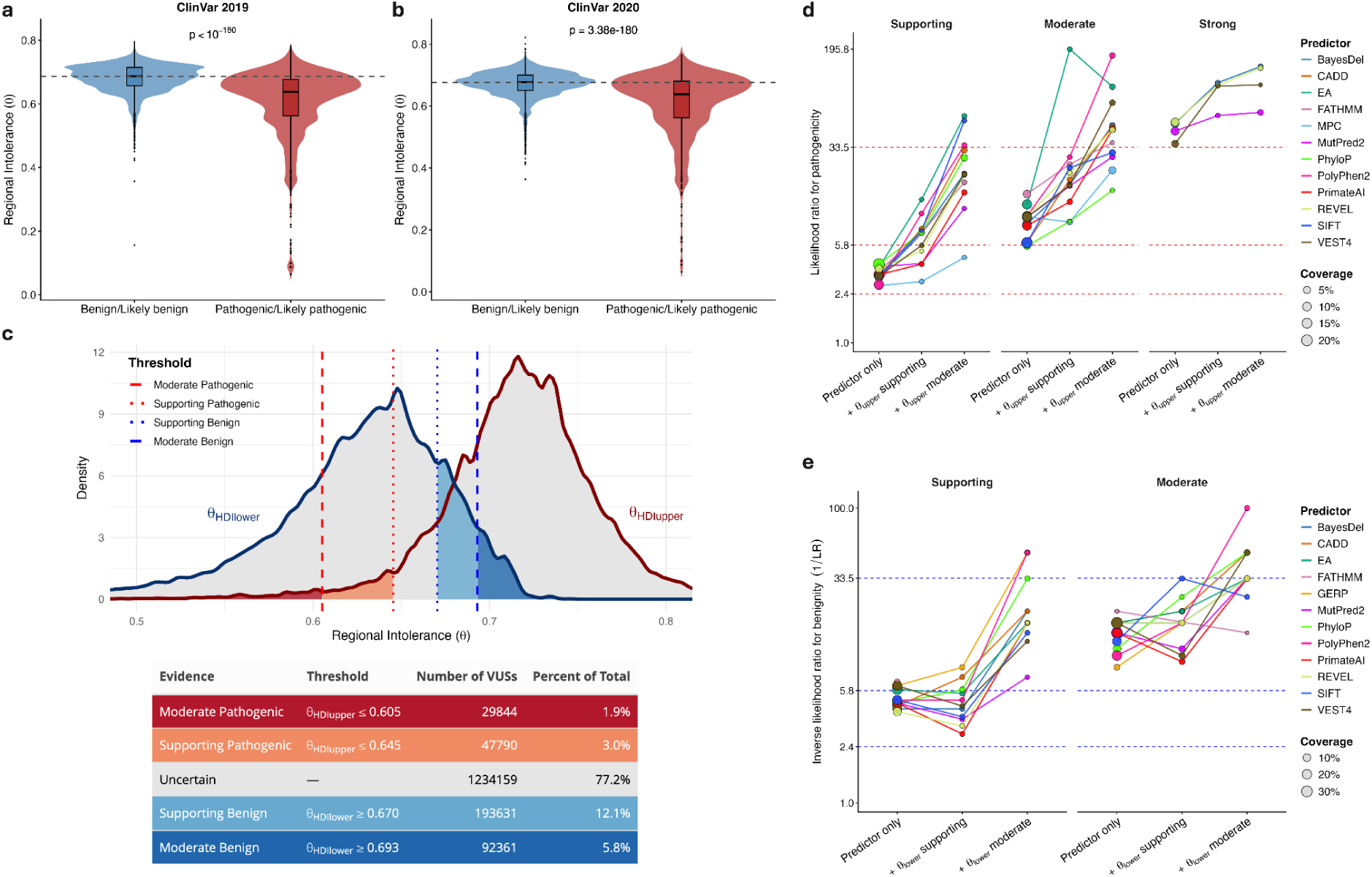
Calibration of PRIME Estimates for Clinical Variant Interpretation. **a**, Distribution of regional intolerance, the posterior mode of θ, for benign and pathogenic variants in the ClinVar 2019 calibration dataset. **b**, Distribution of regional intolerance for benign and pathogenic variants in the ClinVar 2020 validation dataset. Above p-values are from one-sided Mann-Whitney tests. **c**, Distribution of 95% HDI lower and upper bounds among ClinVar VUSs with corresponding VUS classification table below. **d**, Positive likelihood ratios (TPR/FPR) for variant-level predictors across evidence thresholds of pathogenicity, and their change when combined with regional intolerance (θ_HDIupper_) evidence intervals in the ClinVar 2020 validation set. Points show the likelihood ratio for the predictor alone or in conjunction with θ_HDIupper_ supporting or moderate evidence; point size reflects the proportion of variants meeting the interval criterion in the total set of variants with predictions (coverage). Dashed lines represent thresholds of likelihood ratios corresponding to levels of pathogenic evidential strength, from supporting (bottom) to strong (top). **e**, Inverse likelihood ratios 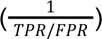 for variant-level predictors across evidence thresholds of benignity, and their change when combined with regional intolerance (θ_HDIlower_) evidence intervals in the ClinVar 2020 validation set. Points show the likelihood ratio for the predictor alone or in conjunction with θ_HDIlower_ supporting or moderate evidence; point size reflects coverage. Dashed lines represent thresholds of likelihood ratios corresponding to levels of benign evidential strength, from supporting (bottom) to strong (top).

Using calibrated PRIME intervals of regional intolerance, missense VUSs present in more recent ClinVar data can be assigned up to moderate evidence of pathogenicity or benignity (Methods: Calibration of PRIME Estimates for Clinical Variant Interpretation). Overall, 363,626/1,597,785 (22.8%) of VUSs fall within genomic regions where θ_HDIlower_ or θ_HDIupper_ meets at least supporting evidence for either classification (Fig. 6c), demonstrating the utility of PRIME in aiding the reclassification of a substantial fraction of currently unresolved missense variants.

It is highly encouraging that a regional annotation derived solely from standing population variation can discriminate pathogenic and benign variants to this degree. However, we acknowledge that this calibration approach was designed for variant-level annotations, and thus may undercall the utility of our model. As such, we sought to determine if PRIME provides complementary information when used in conjunction with variant-level predictors by improving their ability to discriminate between pathogenic and benign mutations, similar to what has been demonstrated for RVIS^1^ and PolyPhen-2^73^. We thus examined how further stratifying variants by calibrated PRIME estimates within previously established predictor-defined evidence intervals^88^ affects their likelihood ratios in the independent ClinVar 2020 validation set (Methods: Calibration of PRIME Estimates for Clinical Variant Interpretation). This approach is conceptually analogous to the consensus-based predictor framework described in Pejaver et al., which combined multiple in silico tools using developer recommended thresholds to approximate their use in certain clinical laboratories. For pathogenicity, further stratifying predictor-defined intervals by supporting regional intolerance evidence resulted in a mean 2.56-fold increase in likelihood ratios across all evidence strengths, increasing to 6.26-fold with moderate intolerance evidence (Fig. 6d, Supplementary Table 3). For benignity, the corresponding mean increases in inverse likelihood ratios were 1.17-fold and 3.84-fold, respectively (Fig. 6e, Supplementary Table 4). As expected, these improvements were accompanied by reduced coverage, as adding progressively stricter intolerance thresholds classified a smaller proportion of variants than just the predictor-defined intervals alone (Supplementary Tables 5,6). We also note that the magnitude of improvement may depend on the extent to which a predictor already incorporates information related to human population variation. For example, MPC (Missense badness, PolyPhen-2, and Constraint) is a composite predictor^17^ that directly incorporates regional missense constraint to distinguish pathogenic variants, and further stratifying by PRIME estimates would not be expected to have a large effect. Consistent with this expectation, MPC showed the smallest improvement in likelihood ratios. Averaged across MPC pathogenic evidence strengths, the addition of supporting PRIME intolerance evidence produced no increase, while moderate PRIME evidence resulted in only a 1.99-fold increase, the smallest effect observed among all predictors in both settings.

## Methods

### Gene Model

Genome annotation was based on the Ensembl v110 release^22^ (GRCh38/hg38) GTF file (Data Availability). Genes were defined based on their canonical transcript in Ensembl, where exons labeled Consensus CDS (CCDS)^21^ were taken as the protein coding regions of the genome. Only canonical, CCDS transcripts with the “protein_coding” transcript biotype were retained. Gene3D^91^ protein domain boundaries derived from the CATH structural hierarchy^92^ were retrieved using the Ensembl v110 BioMart, where their amino acid coordinates were mapped to genomic coordinates using the proteinToGenome function from the ensembldb R package^93^. These steps left us with two annotations of protein coding regions of the genome: a domain-specific gene model based on Gene3D protein domain boundaries and an exon-specific gene model based on Ensembl exon boundaries.

### Variant Annotation

Variant data was downloaded from gnomAD v4.1^4^ from both the exome chromosome-specific VCF files and the genome chromosome-specific VCF files, allowing us to utilize aggregated variant information from 730,947 exomes and 76,215 genomes in our analysis (Data Availability). VCF files from both data sources were filtered to only contain single nucleotide variants (SNVs) with a “PASS” in the gnomAD filter column and located at loci with >=30x median depth from their respective coverage summary files (Data Availability). The VCF files were further filtered to only contain variants that fell within the range of any of our defined protein coding regions from the gene model described above. The Ensembl v110 Variant Effect Predictor (VEP)^94^ was then used to annotate all remaining variants based on their consequence in the canonical transcript of a gene. Variants were only counted for a region if their VEP consequence affected the gene the region is nested within. The “missense_variant” and “synonymous_variant” annotations were counted as missense and synonymous variant counts in a region, respectively, and “start_lost”, “stop_lost”, “stop_gained”, “splice_acceptor_variant”, and “splice_donor_variant” annotations were counted as loss-of-function. Variants that had VEP consequences in multiple genes were counted separately for regions in each gene as the consequence specific to the gene. Genes lacking any standing variation in the population after our filtering steps were discarded, leaving 18,727 genes with 81,749 domain-based subregions or 189,032 exon-based subregions when fitting the PRIME model.

### Estimating Regional Intolerance with PRIME

A Bayesian hierarchical model (Fig. 1b, Supplementary Fig. 1b) was fit to the obtained variant count data to jointly estimate sub-genic intolerance in every defined protein coding region in the genome. We modeled the variant generating process in each subregion *s* with a binomial distribution:

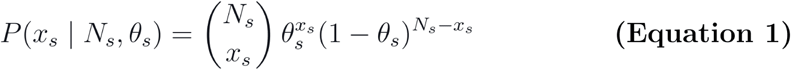

*x_s_* : number of missense variants observed in subregion *s*

*N_s_ :* total number of observed variants (missense, synonymous, and loss-of-function) in subregion *s*

*6S* : probability of a variant being missense in subregion s

More intolerant regions have values of θ closer to 0 in the posterior distribution, as they contain fewer missense variants relative to total variants in the population due to purifying selection. By conditioning on the total number of observed variants, this formulation naturally accounts for regional differences in mutation rate.

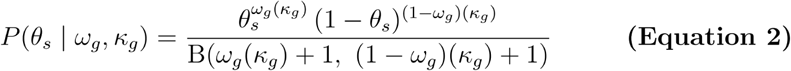

*(Vg* : mode of the beta distribution for gene g

*K_g_* : concentration of the beta distribution for gene g, constrained such that *K_g_ > 2*

*B* : beta function

Regional intolerance is jointly informed by the variants within the region, via the likelihood, and by other regions nested within the same gene, via shrinkage to a shared gene-level estimate. For each gene, we define a beta distribution parameterized by mode and concentration, which serves as a prior for the subregional θ estimates:

This structure induces a shrinkage effect on θ, where regions of the same gene have their estimates pulled towards each other through the prior. In sub-genic regions lacking sufficient variant information, estimates are more strongly informed by the prior. Regions with abundant standing variation are primarily informed by their own data – they converge to the binomial maximum likelihood estimator, *x_s_*/*N_s_*, as the number of total variants increases. The strength of shrinkage within a gene is determined by its concentration parameter κ*_g_*, which is latent in the model. Crucially, this allows for a weaker shrinkage effect in genes with evidence of regional intolerance heterogeneity, and a stronger effect when regional intolerance is mostly homogeneous. The mode of the beta prior, ω*_g_*, is on the same scale as θ, and thus can be interpreted as the probability of a variant being missense in a gene *g*.

207 genes in our final dataset have fewer than 50 total variants. Consequently, their subregions contain just 11.05 and 4.80 variants on average in the domain and exon models respectively, making estimation unstable in many of these regions. To mitigate this, we induced a further dependence in the model across all genes by placing a beta prior on the gene-level modes ω*_g_*:

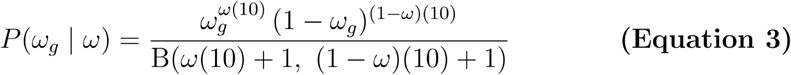

*a>* : mode of the genome-wide beta distribution

This shared prior allowed us to further borrow information across the entire genome. A fixed concentration parameter was chosen to avoid overshrinking gene-level modes, as leaving it latent resulted in impractically high values, even with strong hyperpriors. This was likely due to the parameter being jointly informed by 18,727 genes, in contrast to the gene-level concentration parameters being informed by far fewer subregions.

The PRIME model was fit to the gnomAD variant counts using the rjags R package^95^, which interfaces with JAGS (Just Another Gibbs Sampler)^96^, utilizing 4 parallel chains of 5,000 samples each, totalling 20,000 MCMC samples for all downstream analyses. An adaptation phase of 500 steps followed by burn-in of 500 steps was performed for each chain before posterior samples were retained. The marginal posterior distributions of all latent parameters were summarized by their posterior modes as well as the lower and upper bounds of their 95% highest density intervals.

### Association of Regional Intolerance and ClinVar Missense Pathogenicity

ClinVar^23^ variants were downloaded (Data Availability) and filtered to those annotated as “Pathogenic” or “Likely_pathogenic,” with a “missense_variant” molecular consequence and at least one-star review status. Using the GenomicRanges R package^97^, these variants were counted within all domain-based and exon-based regions based on genomic coordinates, provided they had identical associated gene names. Variants with multiple associated gene names in ClinVar were counted separately for each gene.

To perform the gene-by-gene permutation test assessing how well PRIME estimates predict where pathogenic missense variants fall, an expected number of variants based on neutrality was computed in all domain and exon-based regions. A 7bp sliding window was implemented across all regions to compute the number of expected variants at each locus with a heptamer mutational model, as described in Duan et al^98^. Then, for each region, the expected variants at each locus were summed to get a total number of expected variants in the region. After filtering to genes with at least one ClinVar pathogenic missense variant and at least two subregions, the permutation test could be run on 2,834 genes using domain boundaries and 3,206 genes using exon boundaries. The gene-by-gene permutation procedure described in Hayeck et al.^15^ was then implemented. Briefly, the pathogenic variant counts in regions within a gene were regressed on their PRIME estimates while controlling for the cumulative mutation rate of the region:

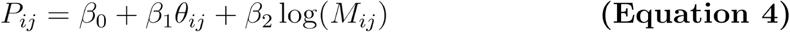

*Pij* : pathogenic variant counts in subregion i of gene j

*dij* : posterior mode of probability of a variant being missense in subregion i of gene j

*Mij :* expected number of variants in subregion i of gene j

The observed slope, β_1_, was recorded for each gene. Then, multinomial sampling was used to permute the pathogenic variant counts in a gene across its subregions proportional to the cumulative number of expected variants. This effectively redistributes the pathogenic variants across the gene as would be expected under neutral mutational burden alone. The regression model above was fit to these 10,000 permuted samples for each gene, creating a null distribution of the slope β_1_ and allowing us to compute p-values by comparing the observed slope to the permuted slopes. Given our expectation of regional intolerance θ and pathogenic counts being negatively correlated, the permutation p-value was calculated as *p* = (G + 1)/10001, where *G* is the number of permutations where 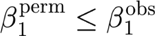. Permutation p-values were adjusted for multiple testing across genes using the Benjamini-Hochberg procedure.

### Association of Regional Intolerance and Experimental DMS Assay Scores

ProteinGym^25^ DMS substitution data was downloaded (Data Availability) and filtered to human samples. The amino acid position of each substitution within each gene was extracted, where multi-mutant variants were ignored. For each assayed variant, the amino acid position, gene name, corresponding DMS score, and binary variable representing whether or not the variant is considered deleterious (DMS_score_bin) was retained. Variants were then assigned domain-based intolerance estimates if the gene names matched and the amino acid coordinate fell within the range of the domain or interdomain region. The same process was repeated for the exon-based intolerance estimates, with the caveat that exon boundaries can fall within a codon of the gene. Thus, DMS variants with amino acid positions that overlapped two exons were assigned to the exon that contained the middle base of the corresponding codon. For both domain and exon-based delineations, the analysis was restricted to genes with at least two subregions, leaving 27 and 30 genes respectively for the follow-up analyses.

A similar permutation procedure to the above ClinVar analysis was performed. For each testable gene, the variant DMS scores were regressed on their corresponding PRIME estimates:

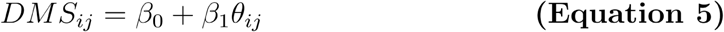

*DMSij* : DMS score of variant i in gene j

*6ij* : posterior mode of regional probability of a variant being missense for variant i of gene j

The observed slope, β_1_, was recorded for each gene. There is no coefficient included for the mutation rate of the region in this analysis as there is no longer a bias in their counts due to mutability. Then, DMS variant scores were shuffled randomly among the regions within a gene without replacement such that each region contained the same number of total variants as in the original assay. The same regression model above was then fit to the reshuffled variant scores and intolerance estimates. This process was performed 10,000 times for each gene, allowing us to compute p-values by comparing the observed β_1_ to the permuted β_1_ values. Given our expectation of regional intolerance θ and DMS scores being positively correlated, the permutation p-value was calculated as *p* = (G + 1)/10001, where *G* is the number of permutations where 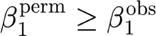. Permutation p-values were adjusted for multiple testing across genes using the Benjamini-Hochberg procedure.

To analyze the enrichment of deleterious variants in intolerant regions of ProteinGym genes, we first quantified each region’s intolerance relative to its gene-wide mean. For each region, relative intolerance was calculated as the regional intolerance estimate from PRIME (*θ_region_*) minus the mean intolerance across all regions within the same gene (*̄θ_gene_*). Negative values indicate regions that are more intolerant than the gene-wide mean, whereas positive values indicate relatively tolerant regions. We then quantified the regional excess of deleterious variants as the difference between the fraction of assayed variants classified as deleterious in the region (DMS_score_bin = 1) and the corresponding fraction across the entire gene. Positive values indicate an excess of deleterious variants relative to the gene-wide baseline, whereas negative values indicate depletion. We then fit weighted regression models for both domain and exon-based regions, regressing this regional difference in the rate of deleterious variants on relative intolerance. Models were weighted by the total number of variants assayed in the region, such that regions with greater variant coverage contributed more strongly to the estimated association. The resulting regression coefficients quantify the relationship between a region’s intolerance relative to its gene-wide mean and its deviation from its gene’s baseline rate of deleterious variants. Negative coefficients indicate that regions more intolerant than their gene-wide mean contain a higher-than-expected fraction of deleterious variants.

### Enrichment Testing of Gene Sets in ClinVar Permutation Test

To test for enrichment of disease gene sets in the set of significant genes from the previous ClinVar permutation test, the OMIM database^27^ (Data Availability) was first queried (Supplementary Methods: Querying the OMIM Database) for seven sets of genes with different disease contexts. The gene sets were defined as such: any gene associated with a Mendelian disorder (“All OMIM”), an autosomal dominant disorder (“Autosomal Dominant”), an autosomal recessive disorder (“Autosomal Recessive”), de novo pathogenic variants (“De Novo”), dominant-negative disease mechanism (“Dominant-Negative”), gain-of-function disease mechanism (“Gain-of-Function”), or haploinsufficiency (“Haploinsufficient”). For each of the seven disease-associated gene sets, a one-sided Fisher’s exact test was used to assess enrichment among genes identified as significant by the ClinVar permutation test based on their unadjusted p-values, relative to the full set of testable genes.

Then, a similar enrichment analysis was performed for the full HGNC hierarchy of gene groups (Data Availability). Broader parent groups of genes encompass more specific child groups, allowing genes to belong to multiple groups across different levels of the hierarchy, which is structured as a directed acyclic graph (DAG). Given there are over 2100 gene groups in this database, a hierarchical gatekeeping procedure^30,31^ was used to control Type I error when testing for their enrichment in the set of significant genes from the ClinVar permutation analysis. First, the root nodes of the DAG were identified, which are the broadest, most encompassing gene groups in the hierarchy. Then, a one-sided Fisher’s exact test was performed for each of these root node gene sets, assuming that all testable genes in the gene set were significant in the ClinVar analysis. This was done to find the minimum possible p-value that gene group could have from the enrichment test. These p-values were subsequently adjusted for multiple testing by the Holm procedure^99^. If the corrected p-values did not reach significance at ɑ = 0.05, the gene group was discarded. For the remaining root-level gene groups, enrichment was assessed using a one-sided Fisher’s exact test comparing the observed number of significant genes (based on unadjusted p-values from the ClinVar permutation tests) within each gene group to the number expected given the overall proportion of significant genes among all testable genes, analogous to the above OMIM enrichment analysis. Only root nodes with Holm-adjusted p-values below ɑ = 0.05 were retained. A hierarchical gatekeeping procedure was subsequently applied, in which only the child nodes of significant parent nodes were tested. At each level of the hierarchy, Holm’s procedure was applied to the p-values of all sibling nodes, and testing proceeded recursively only for child nodes whose parent remained significant after Holm correction. This approach controlled the family-wise error rate across the hierarchy while maintaining power to detect enrichment in more specific gene groups.

### The Weakest Link of Genes

The weakest link statistic for each gene *g* was computed by evaluating, at each MCMC iteration *t*, the minimum intolerance estimate across all *N* subregions, 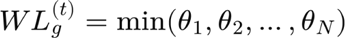, thereby capturing the most intolerant subregion within the gene for each posterior draw. Each gene thus has a posterior distribution of *WL* values, which is then summarized by the upper bound of its 95% HDI, resulting in a conservative estimate of its weakest link. This statistic was computed separately for each gene with domain-based and exon-based subregion definitions. For the AUROC comparison across OMIM gene sets, both statistics were compared to the minimum MCR^17^ observed/expected value for the gene (Data Availability), the missense Z-score^2^ and missense observed/expected upper bound fraction (MOEUF^4^) for the canonical transcripts (Data Availability), RVIS^1^ recomputed with the same variant counts as PRIME, and the upper bound of the 95% HDI of ω*_g_* (Fig. 1b) from PRIME (gene-level intolerance).

To quantify the relative strength of localized intolerance in non-autosomal dominant (non-AD) disease genes, we defined *P_Mendeiun_* as the posterior probability that the weakest link of a given gene is more intolerant than that of a known autosomal dominant (AD) disease gene. For each non-AD gene *g*, at each MCMC iteration *t*, this was computed as:

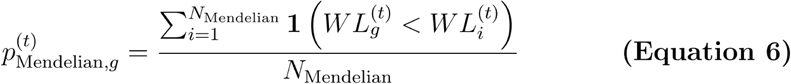

N_Mendalian_ : Number of AD disease genes

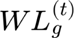 : Weakest link of non-AD disease gene g

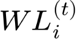 : Weakest link of AD disease gene i

Each non-AD gene thus had a posterior distribution of *P_Mendelian_* values, which was then summarized by the mean.

To investigate the relationship between weakest link intolerance and gene expression, GTEx^38^ median gene-level TPM by tissue (Data Availability) was downloaded and merged with all genes in the analysis. Because expression levels are highly correlated across brain regions, a composite brain expression measure was generated for each gene by taking the median of the gene-level median TPM values across the 13 brain tissue subtypes. This was used for the calculation of the number of tissues a gene is expressed in (Fig. 4f, Supplementary Fig. 5a), making 56 the maximum possible value for a given gene, as well as when computing the median TPM (Supplementary Figs. 4a,5c) and total TPM (Supplementary Figs. 4b,5d) across tissue types for genes. When counting the number of tissues a gene is expressed in, a threshold of TPM >3 was used. The composite brain expression was not used when computing the maximum across tissues (Fig. 4f, Supplementary Fig. 5b) for genes, as that may obscure high expression of a gene in a particular region of the brain. Instead, the maximum was computed using the original tissue-specific values, allowing potential highly localized expression within individual brain regions to contribute to the statistic. For plotting, the maximum, total, and median expression values for genes were transformed as log_10_(TPM + 1) to improve visualization of the highly skewed distributions and accommodate zero values.

### Calibration of PRIME Estimates for Clinical Variant Interpretation

To calibrate the PRIME domain-based estimates for clinical variant interpretation in accordance with ACMG/AMP criteria^89^, the approach outlined in Pejaver et al.^88^ was followed. The provided ClinVar 2019, gnomAD v.2.1, and ClinVar 2020 datasets were first downloaded (Data Availability). All variants found in the mitochondrial DNA were discarded, as only genes on nuclear chromosomes had their regional intolerance estimated with PRIME. Coordinates for the variants were then lifted over from hg19 to hg38 with the UCSC Genome Browser liftover tool^100^.

Using the GenomicRanges package^97^, ClinVar and gnomAD variants were assigned domain-based PRIME estimates by matching Ensembl gene IDs and mapping variant coordinates to domain or interdomain regions. This resulted in 11,535 variants in the ClinVar 2019 set, 355,765 in the gnomAD set, and 8,314 in the ClinVar 2020 set with corresponding PRIME estimates. This included the posterior mode of θ and the lower (θ_HDIlower_) and upper bounds (θ_HDIupper_) of the 95% HDI of θ, which were used for the calibration as they provide conservative estimates in the directions of tolerance and intolerance, respectively.

The ClinVar 2019 and gnomAD v.2.1 datasets were used to calibrate θ_HDIlower_ for benign evidence and θ_HDIupper_ for pathogenic evidence following the procedure outlined by Pejaver et al. Local posterior probabilities were computed for each unique estimate of θ_HDIlower_ and θ_HDIupper_ using adaptive intervals, with a prior probability of 0.0441 for pathogenicity and 1 – 0.0441 (0.9559) for benignity. Each interval was required to contain at least 100 ClinVar variants and at least 3% of gnomAD rare variants to compute posterior probabilities of pathogenicity or benignity at a given estimate. To compute the intervals of θ_HDIlower_ and θ_HDIupper_ corresponding to different levels of evidential support, 10,000 bootstrapped calibrations were performed. Briefly, variants in the ClinVar 2019 and gnomAD v.2.1 datasets were resampled with replacement 10,000 times to generate bootstrap replicates for calibration. This allowed us to compute one-sided 95% confidence bounds of the estimated local likelihood ratio at given values of θ_HDIlower_ or θ_HDIupper_, which were used to determine the final, optimal thresholds for benignity and pathogenicity.

To extract VUSs from a more recent set of ClinVar variants (Data Availability), variants were filtered to those annotated with “Uncertain_significance” clinical significance, a “missense_variant” molecular consequence, and at least one-star review status. VUSs were then assigned domain-based PRIME estimates by matching gene names and mapping variant coordinates to the corresponding domain or interdomain regions. Variants annotated to multiple genes in ClinVar were evaluated separately for each matching gene interpretation. Thresholds computed from the one-sided 95% confidence bounds in the previous step were used to assign pathogenic or benign evidence to variants.

To compare likelihood ratios of using predictor-defined intervals alone to using them with calibrated PRIME intervals, the independent ClinVar 2020 dataset was used. Evidence thresholds of predictors corresponding to ACMG/AMP evidence strengths were obtained from Table 2 of Pejaver et al. and used to assign variants to benign and pathogenic categories based on their predictor scores. Likelihood ratios were computed using Equation 10 in Pejaver et al. for the predictor-defined intervals alone, predictor intervals with supporting PRIME evidence of intolerance, and predictor intervals with moderate PRIME evidence of intolerance. Values were calculated based on the set of variants for a predictor in the ClinVar 2020 set that had both a non-missing value for that predictor and a non-missing value for PRIME. Coverage was similarly computed by taking the percentage of variants that the corresponding interval contained among the set of variants with non-missing values for both.

## Discussion

In this work, we present PRIME, a probabilistic framework for estimating regional missense intolerance across the human genome. Our results demonstrate that modeling intolerance at the level of biologically meaningful subregions can reveal patterns that are not apparent from gene-level analyses alone. These regional estimates provide a foundation for a range of downstream applications, from disease gene discovery to the exploration of structure-function relationships within proteins.

An example of just such a downstream analysis is our inference on each gene’s weakest link. We show that the weakest link can identify genes whose intolerance patterns resemble those of established Mendelian disease genes, including paralogous genes with known pathogenic variants. Notably, of the 634 genes in our dataset with either a domain or exon-derived p_Mendelian_ ≥ 0.8 and an available missense Z-score, 200 (31.5%) fall below the commonly used constraint threshold^2^ of Z = 3.09 (Supplementary Data 8). This highlights many instances where a gene-level metric can dilute signal in highly intolerant regions by aggregating variants across the entire gene. By instead focusing on the weakest link, we provide a robust inferential framework for prioritizing genes based on their most intolerant functional regions, whose disease relevance may otherwise be overlooked by gene-level methods. The need for such an approach in disease gene mapping is reinforced by the observation that many disease genes show a distinct pattern of pathogenic variants clustering in particular regions^5–13^. In some cases, genes that are prioritized by intolerance-based methods may lack recognized Mendelian phenotypes because damaging variants are incompatible with survival and therefore rarely observed. In others, they may underlie milder or incompletely characterized phenotypes that have not yet been linked to specific genes. By focusing on the most missense-depleted subregion within each gene, the weakest link framework provides a powerful approach for identifying genes with Mendelian-like intolerance across a spectrum of phenotypic severity, facilitating their prioritization and interpretation.

A major advantage of PRIME lies in the interpretability of its estimates and in their inference on biologically meaningful units such as protein domains and exons. Unlike approaches that define regions strictly from patterns in the observed data (CCR^16^, MCR^17^), these annotations have direct functional relevance, allowing results to be interpreted in a biological context. As shown in our analysis of selection on domains in gene families (Fig. 5), this allows comparisons to be made within and across genes to explore how their modular protein architecture shapes the distribution of intolerance. These patterns can be highly conserved across paralogs or diverge alongside functional specialization, as observed with the CERS homeobox class (Fig. 5c). Interrogating these broader patterns of selection across gene families may provide insight into the evolution of human proteins and their domains.

Furthermore, the ability to compare intolerance across evolutionarily related proteins may be particularly valuable in a clinical context, where functional redundancy and shared molecular structure can influence disease phenotypes. Consistent with this idea, monogenic disease gene paralogs have been shown to exhibit greater functional similarity than paralogs of non-disease genes^101^. There is also growing evidence that incorporating information from gene families can improve variant interpretation^102,103^. In this respect, PRIME may offer a targeted approach for identifying the domains that are most functionally critical within a gene family and therefore most likely to contribute to disease when disrupted.

We show that PRIME, as a calibrated sub-genic metric, can achieve up to moderate evidence under ACMG/AMP guidelines^89^ for discriminating pathogenic and benign missense variants. Unlike variant-level predictors, PRIME provides region-level annotations across the genome, enabling the identification of subregions intolerant to missense mutation. As such, it offers a framework for prioritizing candidate subregions for targeted functional interrogation, helping focus experimental effort on likely mutational hotspots and coldspots when whole-gene assays are not feasible. This is in fact how many functional assays are performed in practice, where key functional regions are selectively mutated rather than uniformly assaying variants across the entire gene^104–107^. These regions are typically defined by annotated protein domains or prior evidence of disease involvement. However, while domains provide a useful proxy for functional importance, they can serve diverse roles within a protein, and not all are equally sensitive to perturbation. Moreover, selecting regions based on prior disease associations is not feasible for genes with limited existing information. Calibrated regional intolerance estimates from PRIME provide a robust and unbiased framework for identifying functionally important regions - including annotated domains - based on signatures of purifying selection in the population.

We demonstrate how intersecting our calibrated regional intolerance estimates with variant-level predictors can lead to more accurate pathogenicity discrimination, albeit with reduced coverage of variants. This tradeoff is likely the result of applying independently calibrated predictor and intolerance intervals for classification. A more integrated approach that we plan to explore in the future is jointly calibrating such predictors with regional intolerance. Local positive likelihood ratios defined by Equation 6 from Pejaver et al.^88^ could thus be computed while simultaneously considering scores *s* from a continuous variant-level predictor and PRIME estimates *θ*, i.e., modifying *lr^+^*(*s*) to *lr^+^*(*s,θ*). We believe this joint framework will better capture the relationship between variant-level deleteriousness and the fitness consequence of that variant in humans, both of which contribute to disease pathogenicity.

The PRIME model is highly flexible and can be extended to explicitly model additional factors that may affect regional intolerance. In our analysis of gene families, we demonstrate how conserved domains across genes often show similar levels of missense intolerance, likely due to shared structure and function. It is therefore natural to fit a version of PRIME that incorporates random effects for protein domains, allowing information to be shared across them. Domains would then have their intolerance informed not only by other regions within the same gene, but also by homologous domains encoded across the genome, enabling partial pooling across structurally and functionally related protein elements. Previous work has leveraged homologous domains across proteins in a similar manner, either for modeling constraint at a finer resolution^19^ or improving calibration of variant effect predictors^108^. Within PRIME, this can even be implemented in a hierarchical framework using our Gene3D^91^ domain annotations, which are organized according to the CATH structural hierarchy^92^. Top-level structural features, such as secondary structure class, down to more specific functional properties captured at the homologous superfamily level, can be explicitly modeled. We anticipate this approach having two main advantages. First, the borrowing of information across similar domains should allow for more accurate and stable intolerance estimation, particularly for smaller units that are observed across many genes. Second, by incorporating random effects across multiple levels of the CATH structural hierarchy within the probabilistic framework of PRIME, we enable direct comparisons of structural features at varying levels of resolution. At a broad scale, this would allow us to test questions such as whether ɑ-helices are more intolerant to missense variation than β-sheets across the genome. At finer resolution, it enables more specific comparisons, such as whether classical C2H2 zinc finger motifs are more intolerant than other classes of zinc finger domains. As demonstrated in this study, these comparisons would be both interpretable and highly robust, as they propagate uncertainty through the full joint posterior distribution.

Our framework can also be extended to model sub-genic intolerance in all protein-coding transcripts in the genome, moving beyond a focus on canonical isoforms. Previous work has suggested that alternative isoforms of the same gene often function as distinct proteins rather than minor variants of one another, and that this functional divergence should be considered when evaluating their contributions to disease pathogenesis^109^. Failure to account for these alternative transcripts can lead to missed or misinterpreted variants in genetic studies^110^. Modeling regional intolerance at a transcript-specific level can help identify regions most sensitive to perturbation within individual protein isoforms, allowing for more targeted variant interpretation within genes. This extension of the PRIME model fits naturally into its hierarchical structure. The canonical isoform of a gene is typically the longest, and variant information may be sparse in the shorter alternative transcripts, making sub-genic inference of intolerance challenging. Because transcripts are naturally grouped by gene, the model can borrow information both across transcripts within the same gene and across regions within each transcript, thus stabilizing these otherwise noisy estimates. This extension would also allow regions across all transcripts of a gene to be considered when computing its weakest link, potentially revealing disease-relevant regions outside of the canonical isoform.

In summary, PRIME establishes an interpretable and flexible framework to model regional missense intolerance. As shown, this approach can aid in various applications, including prioritizing genes and functionally essential protein domains for disease discovery and clinical variant interpretation. With the continued growth of data available from population sequencing studies, the utility of PRIME in these applications will only further increase.

## Data Availability

### Genome Annotation

GTF gene annotation file was downloaded from the Ensembl FTP site for release 110 (https://ftp.ensembl.org/pub/release-110/gtf/homo_sapiens/Homo_sapiens.GRCh38.110.gtf.gz). **Population Variants**

Exome and genome VCF files with their respective coverage files were downloaded from gnomAD v.4.1.0 (https://gnomad.broadinstitute.org/downloads).

### ClinVar Variants

The ClinVar VCF file used for the permutation tests and VUS analysis can be retrieved from the FTP site (https://ftp.ncbi.nlm.nih.gov/pub/clinvar/vcf_GRCh38/archive_2.0/2025/clinvar_20250608.vcf.gz).

### ProteinGym Assays

ProteinGym DMS assay substitution data was downloaded from their website (https://proteingym.org/download; accessed 2026-02-18).

### OMIM Disease Gene Sets

Disease gene sets were obtained from the OMIM database (https://www.omim.org; accessed 2026-02-12). The full genemap (genemap2.txt) was obtained from the downloads page after registration (https://www.omim.org/downloads).

### HGNC Gene Groups

HGNC gene groups were retrieved from the HGNC database (https://www.genenames.org; accessed 2026-02-17). Hierarchical analysis performed using relevant data (family.csv; hierarchy.csv; hierarchy_closure.csv) from the downloads page (https://www.genenames.org/download/gene-groups).

### Missense Constraint Metrics

Missense constraint regions (MCRs) were obtained from the Google cloud (gs://gcp-public-data--gnomad/papers/2026-rmc; accessed 2026-08-28).

Missense Z-scores and missense observed/expected upper bound fractions (MOEUF) were obtained from the constraint metrics TSV file from the gnomAD v.4.1.1 website (https://gnomad.broadinstitute.org/downloads; accessed 2026-04-15).

### GTEx Tissue Expression

GTEx v10 bulk tissue expression data (GTEx_Analysis_v10_RNASeQCv2.4.2_gene_median_tpm.gct.gz) was downloaded from their portal (https://gtexportal.org/home/downloads/adult-gtex/bulk_tissue_expression).

### ACMG/AMP Calibration Datasets

The ClinVar 2019 (Data S1), gnomAD v.2.0 (Data S2), and ClinVar 2020 (Data S3) datasets were downloaded from the Supplemental information provided in the manuscript (https://pmc.ncbi.nlm.nih.gov/articles/PMC9748256).

## Code Availability

Code used to fit the PRIME model to the raw variant counts and generate summary statistics from the resulting MCMC samples is available online (https://github.com/Costa-Stavrianidis/PRIME.git).

## Acknowledgements

Research reported in this publication was supported by the National Human Genome Research Institute (NHGRI) of the National Institutes of Health (NIH) under award number 5U01-HG011967. Content is solely the responsibility of the authors.

## Notes

https://github.com/Costa-Stavrianidis/PRIME.git

### Competing Interest Statement

The authors have declared no competing interest.

## References

1. Petrovski, S., Wang, Q., Heinzen, E. L., Allen, A. S. & Goldstein, D. B. Genic Intolerance to Functional Variation and the Interpretation of Personal Genomes. PLoS Genet. 9, e1003709 (2013).

2. Samocha, K. E. et al. A framework for the interpretation of de novo mutation in human disease. Nat. Genet. 46, 944–950 (2014).

3. Lek, M. et al. Analysis of protein-coding genetic variation in 60,706 humans. Nature 536, 285–291 (2016).

4. Karczewski, K. J. et al. The mutational constraint spectrum quantified from variation in 141,456 humans. Nature 581, 434–443 (2020).

5. Goldstein, D. B. et al. Sequencing studies in human genetics: design and interpretation. Nat. Rev. Genet. 14, 460–470 (2013).

6. Eilbeck, K., Quinlan, A. & Yandell, M. Settling the score: variant prioritization and Mendelian disease. Nat. Rev. Genet. 18, 599–612 (2017).

7. Dietz, H. C., Saraiva, J. M., Pyeritz, R. E., Cutting, G. R. & Francomano, C. A. Clustering of fibrillin (FBN1) missense mutations in Marfan syndrome patients at cysteine residues in EGF-like domains. Hum. Mutat. 1, 366–374 (1992).

8. Talbot, K. et al. Missense mutation clustering in the survival motor neuron gene: a role for a conserved tyrosine and glycine rich region of the protein in RNA metabolism? Hum. Mol. Genet. 6, 497–500 (1997).

9. Wang, C. M. et al. Identification of 13 novel NLRP7 mutations in 20 families with recurrent hydatidiform mole; missense mutations cluster in the leucine-rich region. J. Med. Genet. 46, 569–575 (2009).

10. Kamburov, A. et al. Comprehensive assessment of cancer missense mutation clustering in protein structures. Proc. Natl. Acad. Sci. U. S. A. 112, E5486–5495 (2015).

11. Turner, T. N. et al. Proteins linked to autosomal dominant and autosomal recessive disorders harbor characteristic rare missense mutation distribution patterns. Hum. Mol. Genet. 24, 5995–6002 (2015).

12. Homburger, J. R. et al. Multidimensional structure-function relationships in human β-cardiac myosin from population-scale genetic variation. Proc. Natl. Acad. Sci. U. S. A. 113, 6701–6706 (2016).

13. Quinodoz, M. et al. Analysis of missense variants in the human genome reveals widespread gene-specific clustering and improves prediction of pathogenicity. Am. J. Hum. Genet. 109, 457–470 (2022).

14. Gussow, A. B., Petrovski, S., Wang, Q., Allen, A. S. & Goldstein, D. B. The intolerance to functional genetic variation of protein domains predicts the localization of pathogenic mutations within genes. Genome Biol. 17, 9 (2016).

15. Hayeck, T. J. et al. Improved Pathogenic Variant Localization via a Hierarchical Model of Sub-regional Intolerance. Am. J. Hum. Genet. 104, 299–309 (2019).

16. Havrilla, J. M., Pedersen, B. S., Layer, R. M. & Quinlan, A. R. A map of constrained coding regions in the human genome. Nat. Genet. 51, 88–95 (2019).

17. Wang, L. et al. The landscape of regional missense mutational intolerance quantified from 730,947 exomes. 2024.04.11.588920 Preprint at 10.1101/2024.04.11.588920 (2026).

18. Traynelis, J. et al. Optimizing genomic medicine in epilepsy through a gene-customized approach to missense variant interpretation. Genome Res. 27, 1715–1729 (2017).

19. Zhang, X. et al. Genetic constraint at single amino acid resolution in protein domains improves missense variant prioritisation and gene discovery. Genome Med. 16, 88 (2024).

20. Li, B., Roden, D. M. & Capra, J. A. The 3D mutational constraint on amino acid sites in the human proteome. Nat. Commun. 13, 3273 (2022).

21. Pujar, S. et al. Consensus coding sequence (CCDS) database: a standardized set of human and mouse protein-coding regions supported by expert curation. Nucleic Acids Res. 46, D221–D228 (2018).

22. Martin, F. J. et al. Ensembl 2023. Nucleic Acids Res. 51, D933–D941 (2023).

23. Landrum, M. J. et al. ClinVar: public archive of relationships among sequence variation and human phenotype. Nucleic Acids Res. 42, D980–D985 (2014).

24. Haghshenas, S. et al. Menke-Hennekam syndrome; delineation of domain-specific subtypes with distinct clinical and DNA methylation profiles. Hum. Genet. Genomics Adv. 5, 100287 (2024).

25. Notin, P. et al. ProteinGym: Large-Scale Benchmarks for Protein Design and Fitness Prediction. Preprint at 10.1101/2023.12.07.570727 (2023).

26. Kroker, A. J. & Bruning, J. B. Review of the Structural and Dynamic Mechanisms of PPARγ Partial Agonism. PPAR Res. 2015, 816856 (2015).

27. Hamosh, A., Scott, A. F., Amberger, J. S., Bocchini, C. A. & McKusick, V. A. Online Mendelian Inheritance in Man (OMIM), a knowledgebase of human genes and genetic disorders. Nucleic Acids Res. 33, D514–517 (2005).

28. Balick, D. J., Jordan, D. M., Sunyaev, S. & Do, R. Overcoming constraints on the detection of recessive selection in human genes from population frequency data. Am. J. Hum. Genet. 109, 33–49 (2022).

29. Seal, R. L. et al. Genenames.org: the HGNC and PGNC resources in 2026. Nucleic Acids Res. 54, D1098–D1107 (2026).

30. Westfall, P. H. & Krishen, A. Optimally weighted, fixed sequence and gatekeeper multiple testing procedures. J. Stat. Plan. Inference 99, 25–40 (2001).

31. Dmitrienko, A., Wiens, B. L., Tamhane, A. C. & Wang, X. Tree-structured gatekeeping tests in clinical trials with hierarchically ordered multiple objectives. Stat. Med. 26, 2465–2478 (2007).

32. Saleem, R. A., Banerjee-Basu, S., Berry, F. B., Baxevanis, A. D. & Walter, M. A. Structural and functional analyses of disease-causing missense mutations in the forkhead domain of FOXC1. Hum. Mol. Genet. 12, 2993–3005 (2003).

33. Reis, L. M. et al. Axenfeld-Rieger syndrome: more than meets the eye. J. Med. Genet. 60, 368–379 (2023).

34. Zhou, L. et al. Genotype-phenotype association of PITX2 and FOXC1 in Axenfeld-Rieger syndrome. Exp. Eye Res. 226, 109307 (2023).

35. Yatsenko, S. A. et al. The Human Intolerome: A curated database to prioritize genomic variants in stillbirth, pregnancy loss, and neonatal death. Genet. Med. Off. J. Am. Coll. Med. Genet. 28, 102546 (2026).

36. Dickinson, M. E. et al. High-throughput discovery of novel developmental phenotypes. Nature 537, 508–514 (2016).

37. Wilson, R. et al. International Mouse Phenotyping Consortium Portal: facilitating investigation of gene function and providing insights into human disease. Nucleic Acids Res. 54, D1133–D1142 (2026).

38. Lonsdale, J. et al. The Genotype-Tissue Expression (GTEx) project. Nat. Genet. 45, 580–585 (2013).

39. Buscaglia, G., Northington, K. R., Aiken, J., Hoff, K. J. & Bates, E. A. Bridging the Gap: The Importance of TUBA1A α-Tubulin in Forming Midline Commissures. Front. Cell Dev. Biol. 9, 789438 (2021).

40. Arsenian, S., Weinhold, B., Oelgeschläger, M., Rüther, U. & Nordheim, A. Serum response factor is essential for mesoderm formation during mouse embryogenesis. EMBO J. 17, 6289–6299 (1998).

41. Saccone, S. et al. Origin and Evolution of Genes in Eukaryotes: Mechanisms, Dynamics, and Functional Implications. Genes 16, 702 (2025).

42. Fong, J. H., Geer, L. Y., Panchenko, A. R. & Bryant, S. H. Modeling the Evolution of Protein Domain Architectures Using Maximum Parsimony. J. Mol. Biol. 366, 307–315 (2007).

43. Forslund, K., Pekkari, I. & Sonnhammer, E. L. Domain architecture conservation in orthologs. BMC Bioinformatics 12, 326 (2011).

44. Faulkner, I. E., Pajak, R. Z., Harte, M. K., Glazier, J. D. & Hager, R. Voltage-gated potassium channels as a potential therapeutic target for the treatment of neurological and psychiatric disorders. Front. Cell. Neurosci. 18, 1449151 (2024).

45. Wulff, H., Castle, N. A. & Pardo, L. A. Voltage-gated Potassium Channels as Therapeutic Drug Targets. Nat. Rev. Drug Discov. 8, 982–1001 (2009).

46. Soler-Llavina, G. J., Chang, T.-H. & Swartz, K. J. Functional interactions at the interface between voltage-sensing and pore domains in the Shaker K(v) channel. Neuron 52, 623–634 (2006).

47. Jiang, Y. et al. X-ray structure of a voltage-dependent K+ channel. Nature 423, 33–41 (2003).

48. Lu, Z., Klem, A. M. & Ramu, Y. Ion conduction pore is conserved among potassium channels. Nature 413, 809–813 (2001).

49. Lu, Z., Klem, A. M. & Ramu, Y. Coupling between Voltage Sensors and Activation Gate in Voltage-gated K+ Channels. J. Gen. Physiol. 120, 663–676 (2002).

50. Bocksteins, E. & Snyders, D. J. Electrically silent Kv subunits: their molecular and functional characteristics. Physiology 27, 73–84 (2012).

51. Li, S., Wong, A. H. C. & Liu, F. Ligand-gated ion channel interacting proteins and their role in neuroprotection. Front. Cell. Neurosci. 8, 125 (2014).

52. Nys, M., Kesters, D. & Ulens, C. Structural insights into Cys-loop receptor function and ligand recognition. Biochem. Pharmacol. 86, 1042–1053 (2013).

53. Gielen, M., Thomas, P. & Smart, T. G. The desensitization gate of inhibitory Cys-loop receptors. Nat. Commun. 6, 6829 (2015).

54. Macdonald, R. L., Kang, J.-Q. & Gallagher, M. J. Mutations in GABAA receptor subunits associated with genetic epilepsies. J. Physiol. 588, 1861–1869 (2010).

55. Hernandez, C. C. et al. GABRG2 Variants Associated with Febrile Seizures. Biomolecules 13, 414 (2023).

56. Engel, A. G., Ohno, K. & Sine, S. M. Congenital Myasthenic Syndromes: Recent Advances. Arch. Neurol. 56, 163–167 (1999).

57. Forman, S. A. & Miller, K. W. Anesthetic Sites and Allosteric Mechanisms of Action on Cys-loop Ligand-gated Ion Channels. Can. J. Anaesth. J. Can. Anesth. 58, 191–205 (2011).

58. Zhu, S. et al. Structural and dynamic mechanisms of GABAA receptor modulators with opposing activities. Nat. Commun. 13, 4582 (2022).

59. Lynagh, T. & Pless, S. A. Principles of agonist recognition in Cys-loop receptors. Front. Physiol. 5, 160 (2014).

60. Lewis, D. L. et al. Ectopic gene expression and homeotic transformations in arthropods using recombinant Sindbis viruses. Curr. Biol. CB 9, 1279–1287 (1999).

61. Duverger, O. & Morasso, M. I. Role of homeobox genes in the patterning, specification and differentiation of ectodermal appendages in mammals. J. Cell. Physiol. 216, 337–346 (2008).

62. Hobert, O. & Westphal, H. Functions of LIM-homeobox genes. Trends Genet. TIG 16, 75–83 (2000).

63. Holland, P. W., Booth, H. A. F. & Bruford, E. A. Classification and nomenclature of all human homeobox genes. BMC Biol. 5, 47 (2007).

64. Bürglin, T. R. & Affolter, M. Homeodomain proteins: an update. Chromosoma 125, 497–521 (2016).

65. Li, Z. et al. The RNA-Binding Motif Protein Family in Cancer: Friend or Foe? Front. Oncol. 11, 757135 (2021).

66. Gerstberger, S., Hafner, M., Ascano, M. & Tuschl, T. Evolutionary conservation and expression of human RNA-binding proteins and their role in human genetic disease. Adv. Exp. Med. Biol. 825, 1–55 (2014).

67. Gebauer, F., Schwarzl, T., Valcárcel, J. & Hentze, M. W. RNA-binding proteins in human genetic disease. Nat. Rev. Genet. 22, 185–198 (2021).

68. Busch, A. & Hertel, K. J. EVOLUTION OF SR PROTEIN AND HnRNP SPLICING REGULATORY FACTORS. Wiley Interdiscip. Rev. RNA 3, 1–12 (2012).

69. Ly, J., Cady, S. L., Haug, S., Khalizeva, E. & Cheeseman, I. M. SRSF12 is a primate-specific splicing factor that induces a tissue-specific gene expression program. Mol. Biol. Cell 36, ar138 (2025).

70. Burke, W., Parens, E., Chung, W. K., Berger, S. M. & Appelbaum, P. S. The challenge of genetic variants of uncertain clinical significance: A narrative review. Ann. Intern. Med. 175, 994–1000 (2022).

71. Molotkov, I., Mardis, E. R. & Artomov, M. Making sense of missense: challenges and opportunities in variant pathogenicity prediction. Dis. Model. Mech. 17, dmm052218 (2024).

72. Schmidt, A. et al. Predicting the pathogenicity of missense variants using features derived from AlphaFold2. Bioinformatics 39, btad280 (2023).

73. Adzhubei, I., Jordan, D. M. & Sunyaev, S. R. Predicting Functional Effect of Human Missense Mutations Using PolyPhen-2. Curr. Protoc. Hum. Genet. Editor. Board Jonathan Haines Al 0 7, Unit7.20 (2013).

74. Ng, P. C. & Henikoff, S. Predicting deleterious amino acid substitutions. Genome Res. 11, 863–874 (2001).

75. Kircher, M. et al. A general framework for estimating the relative pathogenicity of human genetic variants. Nat. Genet. 46, 310–315 (2014).

76. Ioannidis, N. M. et al. REVEL: An Ensemble Method for Predicting the Pathogenicity of Rare Missense Variants. Am. J. Hum. Genet. 99, 877–885 (2016).

77. Feng, B.-J. PERCH: A Unified Framework for Disease Gene Prioritization. Hum. Mutat. 38, 243–251 (2017).

78. Pejaver, V. et al. Inferring the molecular and phenotypic impact of amino acid variants with MutPred2. Nat. Commun. 11, 5918 (2020).

79. Katsonis, P. & Lichtarge, O. A formal perturbation equation between genotype and phenotype determines the Evolutionary Action of protein-coding variations on fitness. Genome Res. 24, 2050–2058 (2014).

80. Shihab, H. A. et al. Predicting the functional, molecular, and phenotypic consequences of amino acid substitutions using hidden Markov models. Hum. Mutat. 34, 57–65 (2013).

81. Davydov, E. V. et al. Identifying a high fraction of the human genome to be under selective constraint using GERP++. PLoS Comput. Biol. 6, e1001025 (2010).

82. Pollard, K. S., Hubisz, M. J., Rosenbloom, K. R. & Siepel, A. Detection of nonneutral substitution rates on mammalian phylogenies. Genome Res. 20, 110–121 (2010).

83. Sundaram, L. et al. Predicting the clinical impact of human mutation with deep neural networks. Nat. Genet. 50, 1161–1170 (2018).

84. Carter, H., Douville, C., Stenson, P. D., Cooper, D. N. & Karchin, R. Identifying Mendelian disease genes with the variant effect scoring tool. BMC Genomics 14 **Suppl 3**, S3 (2013).

85. Cheng, J. et al. Accurate proteome-wide missense variant effect prediction with AlphaMissense. Science 381, eadg7492 (2023).

86. Brandes, N., Goldman, G., Wang, C. H., Ye, C. J. & Ntranos, V. Genome-wide prediction of disease variant effects with a deep protein language model. Nat. Genet. 55, 1512–1522 (2023).

87. Tavtigian, S. V. et al. Modeling the ACMG/AMP Variant Classification Guidelines as a Bayesian Classification Framework. Genet. Med. Off. J. Am. Coll. Med. Genet. 20, 1054–1060 (2018).

88. Pejaver, V. et al. Calibration of computational tools for missense variant pathogenicity classification and ClinGen recommendations for PP3/BP4 criteria. Am. J. Hum. Genet. 109, 2163–2177 (2022).

89. Richards, S. et al. Standards and Guidelines for the Interpretation of Sequence Variants: A Joint Consensus Recommendation of the American College of Medical Genetics and Genomics and the Association for Molecular Pathology. Genet. Med. Off. J. Am. Coll. Med. Genet. 17, 405–424 (2015).

90. Harrison, S. M., Biesecker, L. G. & Rehm, H. L. Overview of specifications to the ACMG/AMP variant interpretation guidelines. Curr. Protoc. Hum. Genet. 103, e93 (2019).

91. Lewis, T. E. et al. Gene3D: Extensive prediction of globular domains in proteins. Nucleic Acids Res. 46, D435–D439 (2018).

92. Waman, V. P. et al. CATH v4.4: major expansion of CATH by experimental and predicted structural data. Nucleic Acids Res. 53, D348–D355 (2025).

93. Rainer, J., Gatto, L. & Weichenberger, C. X. ensembldb: an R package to create and use Ensembl-based annotation resources. Bioinformatics 35, 3151–3153 (2019).

94. McLaren, W. et al. The Ensembl Variant Effect Predictor. Genome Biol. 17, 122 (2016).

95. Plummer, M. Rjags: Bayesian Graphical Models Using MCMC. (2025).

96. Plummer, M. JAGS: A program for analysis of Bayesian graphical models using Gibbs sampling. in Proceedings of the 3rd International Workshop on Distributed Statistical Computing (DSC 2003) vol. 124 1–10 (Vienna, Austria, 2003).

97. Lawrence, M. et al. Software for computing and annotating genomic ranges. PLoS Comput. Biol. 9, e1003118 (2013).

98. Duan, Y. et al. Functional Category-Specific Intolerance Reflects Genic Function and Clinical Relevance. 2025.09.24.678298 Preprint at 10.1101/2025.09.24.678298 (2025).

99. Holm, S. A Simple Sequentially Rejective Multiple Test Procedure. Scand. J. Stat. 6, 65–70 (1979).

100. Casper, J. et al. The UCSC Genome Browser database: 2026 update. Nucleic Acids Res. 54, D1331–D1335 (2025).

101. Chen, W.-H., Zhao, X.-M., van Noort, V. & Bork, P. Human monogenic disease genes have frequently functionally redundant paralogs. PLoS Comput. Biol. 9, e1003073 (2013).

102. Lal, D. et al. Gene family information facilitates variant interpretation and identification of disease-associated genes in neurodevelopmental disorders. Genome Med. 12, 28 (2020).

103. Pérez-Palma, E. et al. Identification of pathogenic variant enriched regions across genes and gene families. Genome Res. 30, 62–71 (2020).

104. Findlay, G. M. et al. Accurate classification of BRCA1 variants with saturation genome editing. Nature 562, 217–222 (2018).

105. Sahu, S. et al. Saturation genome editing of 11 codons and exon 13 of BRCA2 coupled with chemotherapeutic drug response accurately determines pathogenicity of variants. PLOS Genet. 19, e1010940 (2023).

106. Beltran, A., Jiang, X., Shen, Y. & Lehner, B. Site-saturation mutagenesis of 500 human protein domains. Nature 637, 885–894 (2025).

107. Funk, J. S. et al. Deep CRISPR mutagenesis characterizes the functional diversity of TP53 mutations. Nat. Genet. 57, 140–153 (2025).

108. Chen, Y. et al. Gene- and domain-aware calibration increases the clinical utility of variant effect predictors. Preprint at 10.64898/2026.02.17.706269 (2026).

109. Yang, X. et al. Widespread expansion of protein interaction capabilities by alternative splicing. Cell 164, 805–817 (2016).

110. Schoch, K. et al. Alternative transcripts in variant interpretation: the potential for missed diagnoses and misdiagnoses. Genet. Med. Off. J. Am. Coll. Med. Genet. 22, 1269–1275 (2020).

