## Supplementary Information for "Probabilistic mapping of sub-genic intolerance reveals functional and disease-critical protein regions"

**Querying the OMIM Database**

To obtain the “All OMIM” (n=4,270 genes) dataset, the full genemap was downloaded and filtered for genes with a phenotype that did not include the terms “resistance”, “cancer”, “somatic”, “susceptibility”, “carcinoma”, and “tumor” or symbols “{“, “[]”, and “?”, as outlined by Petrovski et al.[1]. The “Autosomal Dominant” (n=1,606) and “Autosomal Recessive” (n=2,830) genes were extracted by further filtering the full dataset for phenotypes containing the term “Autosomal dominant” and “Autosomal recessive”, respectively. The “De Novo” (n=1,305), “Dominant-Negative” (n=676), “Gain-of-Function” (n=486), and “Haploinsufficient” (n=277) gene sets were compiled by querying the terms “de novo”, “dominant-negative”, “gain-of-function”, and “haploinsufficiency”, filtering only for gene entries annotated with “*” (known sequence) and “#” (phenotype description and molecular basis known). Records were then further restricted to require “Allelic variants” and a “Gene Map Locus”. The overlap among these gene sets is summarized in Supplementary Table 7.

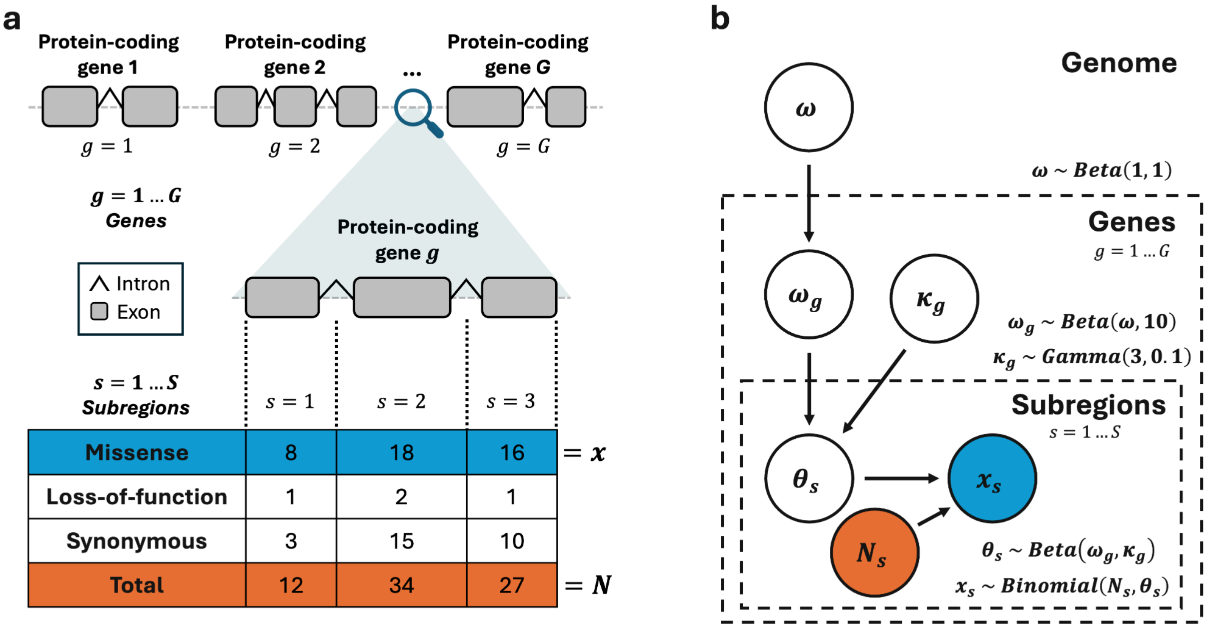


**Supplementary Figure 1: PRIME Framework and Model for Exon-Based Subregions.**
**a**, Workflow to obtain regional variant counts for exons. Protein-coding genes are divided into exons based on Ensembl annotations, and gnomAD exome and genome variants are counted in each region after consequence annotation. **b**, Graphical model of PRIME, fit to obtained variant counts from previous step. Colored nodes denote observed data and unfilled nodes denote latent parameters. PRIME jointly estimates θ, the probability of a variant in the population within an exon being missense. Hyperpriors match those used for the domain-based model in Fig. 1b.

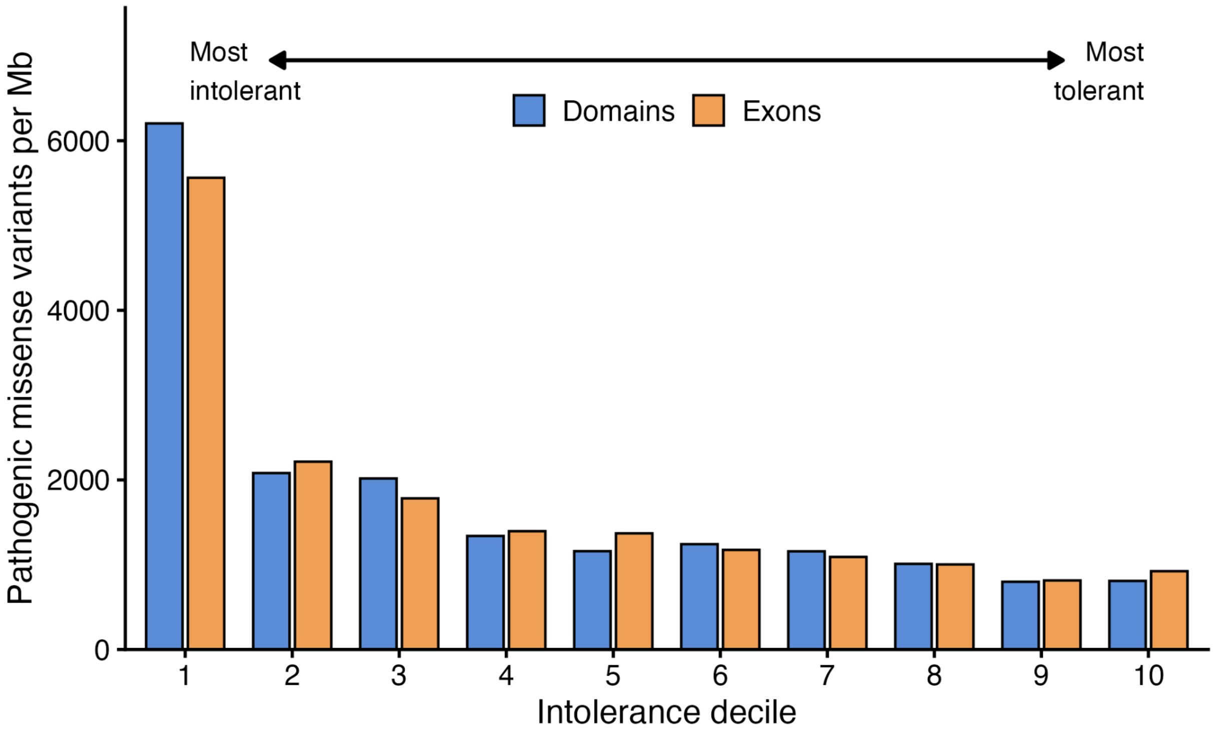


**Supplementary Figure 2: Pathogenic Variants per Megabase by Intolerance Deciles.**
ClinVar pathogenic missense variant counts per megabase for equally sized deciles of intolerance. Intolerant regions demonstrate a higher rate of pathogenic variants relative to more tolerant regions across the genome for both domain and exon-based regions.

| Gene Set | Weakest Link (Domains) | Weakest Link (Exons) | MCR O/E Minimum | Missense Z | MOEUF | RVIS | Omega HDI Upper Bound |
|---|---|---|---|---|---|---|---|
| De Novo | 0.749 [0.733, 0.764] | **0.759 [0.744, 0.774]** | 0.747 [0.733, 0.762] | 0.750 [0.735, 0.766] | 0.740 [0.726, 0.756] | 0.743 [0.727, 0.758] | 0.725 [0.709, 0.740] |
| Gain-of-Function | **0.737 [0.713, 0.759]** | 0.725 [0.700, 0.749] | 0.725 [0.700, 0.750] | 0.726 [0.701, 0.750] | 0.714 [0.691, 0.737] | 0.727 [0.702, 0.753] | 0.730 [0.704, 0.754] |
| Haploinsufficient | 0.727 [0.694, 0.758] | 0.729 [0.696, 0.761] | 0.704 [0.669, 0.737] | 0.671 [0.633, 0.708] | 0.661 [0.626, 0.696] | **0.738 [0.702, 0.771]** | 0.702 [0.666, 0.735] |
| Dominant-Negative | 0.711 [0.690, 0.731] | 0.702 [0.681, 0.723] | 0.708 [0.686, 0.728] | **0.714 [0.691, 0.733]** | 0.704 [0.684, 0.724] | 0.697 [0.674, 0.719] | 0.695 [0.674, 0.716] |
| Autosomal Dominant | **0.710 [0.697, 0.724]** | 0.703 [0.688, 0.716] | 0.704 [0.690, 0.717] | 0.697 [0.682, 0.712] | 0.687 [0.673, 0.702] | 0.707 [0.692, 0.722] | 0.702 [0.688, 0.716] |
| All OMIM | 0.586 [0.576, 0.595] | **0.618 [0.608, 0.627]** | 0.567 [0.557, 0.577] | 0.575 [0.565, 0.585] | 0.570 [0.561, 0.580] | 0.567 [0.557, 0.578] | 0.596 [0.586, 0.606] |
| Autosomal Recessive | 0.494 [0.483, 0.505] | **0.540 [0.528, 0.550]** | 0.464 [0.452, 0.475] | 0.480 [0.469, 0.491] | 0.475 [0.465, 0.485] | 0.471 [0.458, 0.483] | 0.523 [0.512, 0.535] |

**Supplementary Table 1: AUROCs of Intolerance Metrics Discriminating OMIM Disease Gene Sets.** AUROCs of intolerance metrics with 95% confidence intervals in brackets. Confidence intervals were estimated using 2,000 bootstrap resamples of each dataset. Top performing method based on AUROC for a given gene set is bolded.

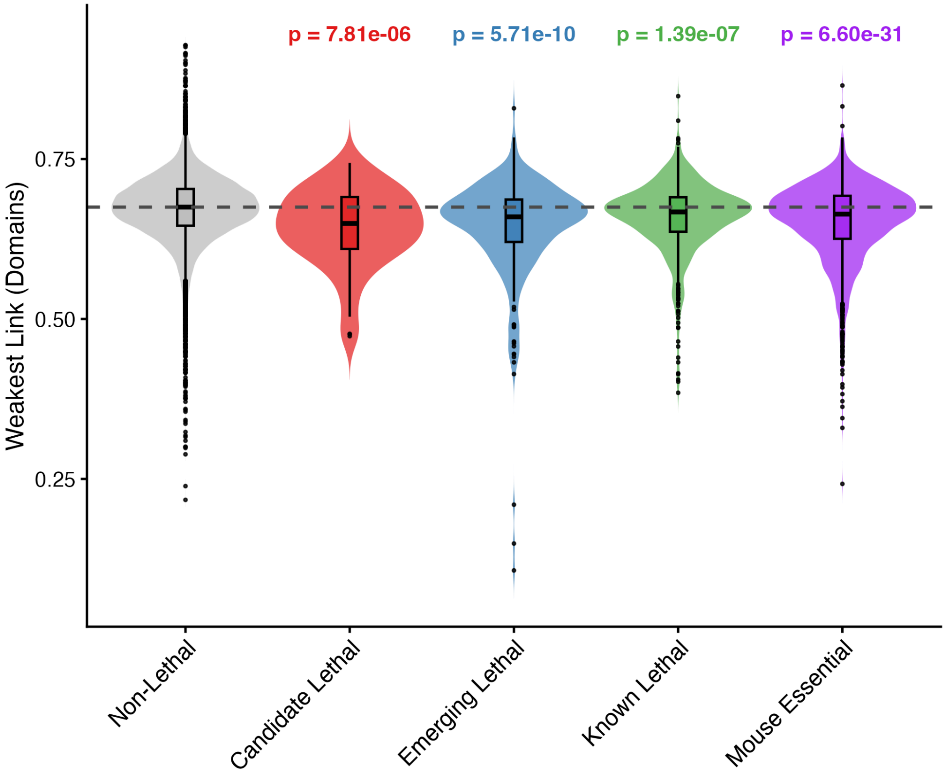


**Supplementary Figure 3: Lethal/Essential Genes Domain-Based Weakest Link Intolerance.** Distribution of the domain-based weakest link statistic for lethal genes with varying levels of evidence and genes essential for development in mice. Above p-values are from one-sided Mann-Whitney tests comparing the distributional differences between the lethal/essential gene sets and the complement of those sets combined (non-lethal genes). Dashed line represents the median weakest link value among non-lethal group.

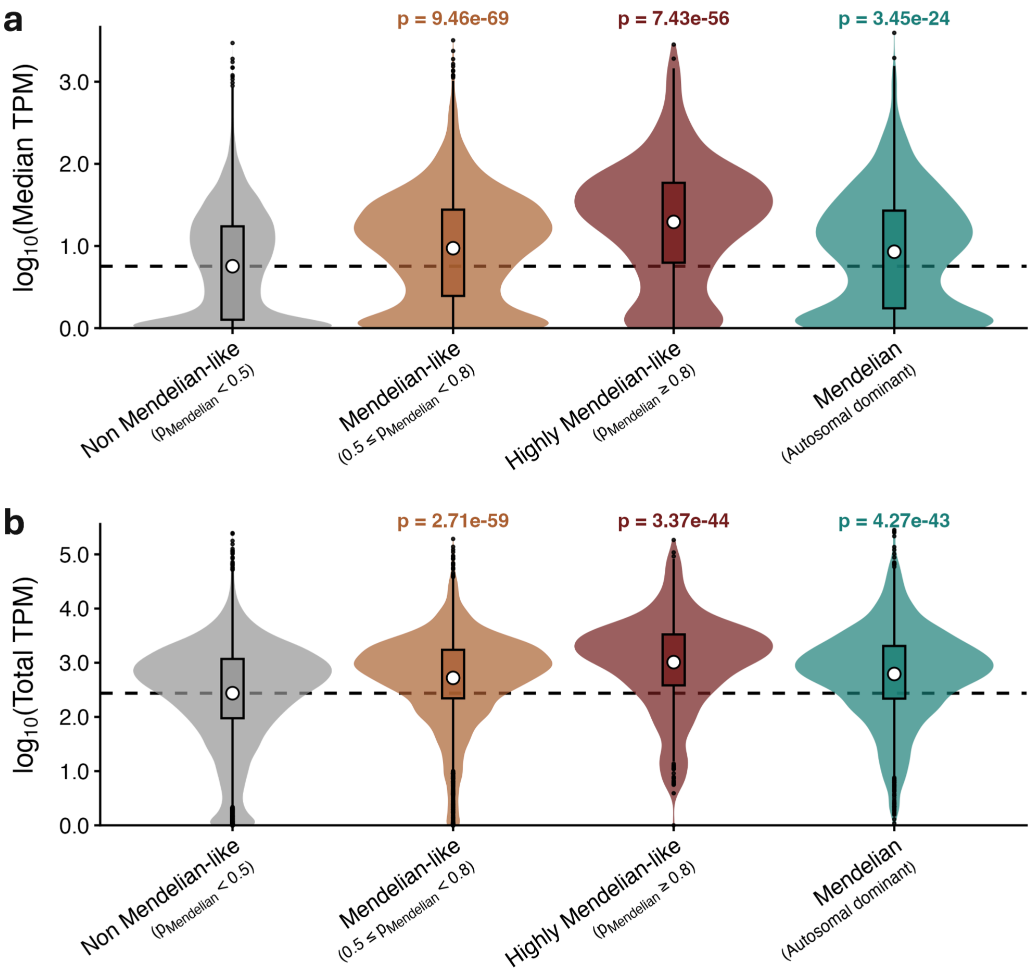


**Supplementary Figure 4: Distribution of Domain-based Mendelian-like and Non Mendelian-like Gene Expression.**

**a,** Mendelian-like genes and known Mendelian disease genes show higher median expression across tissue types compared to non-Mendelian genes. **b,** Mendelian-like genes and known Mendelian disease genes show higher total expression across tissue types compared to non-Mendelian genes. Dashed lines represent the mean values of the non Mendelian-like set.

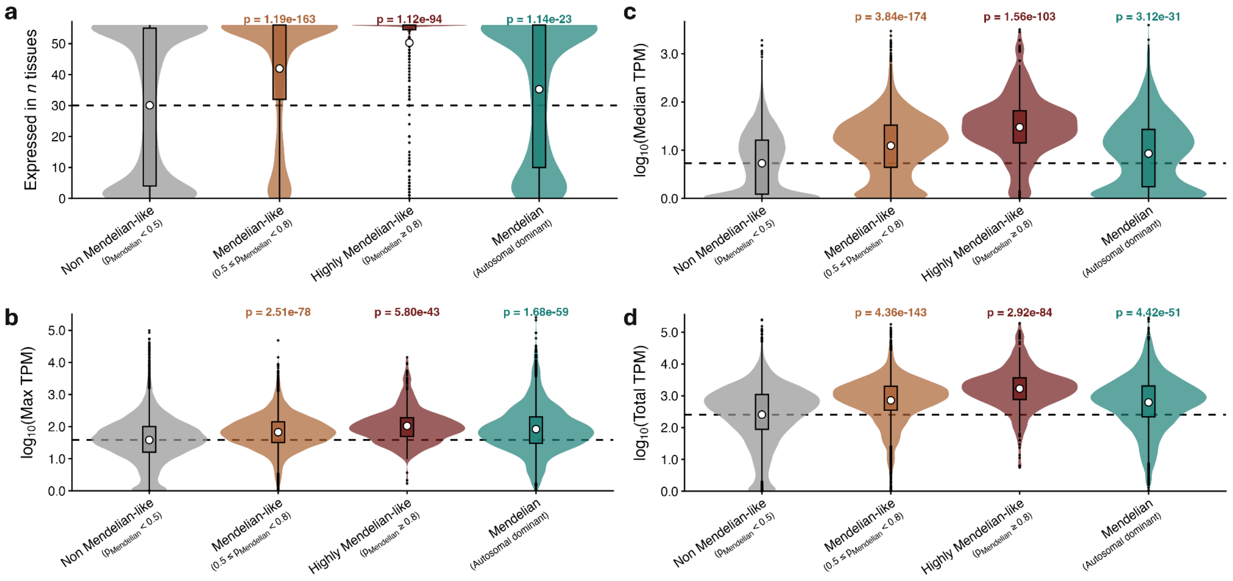


**Supplementary Figure 5: Distribution of Exon-based Mendelian-like and Non Mendelian-like Gene Expression.**

**a,** Number of tissues expressed, **b,** maximum tissue-specific expression, **c,** median expression across tissues, and **d,** total expression across tissues for non-Mendelian, Mendelian-like, and known Mendelian disease genes. Mendelian-like genes and known Mendelian disease genes exhibit higher expression across all four measures. Dashed lines represent the mean values of the non-Mendelian like gene set.

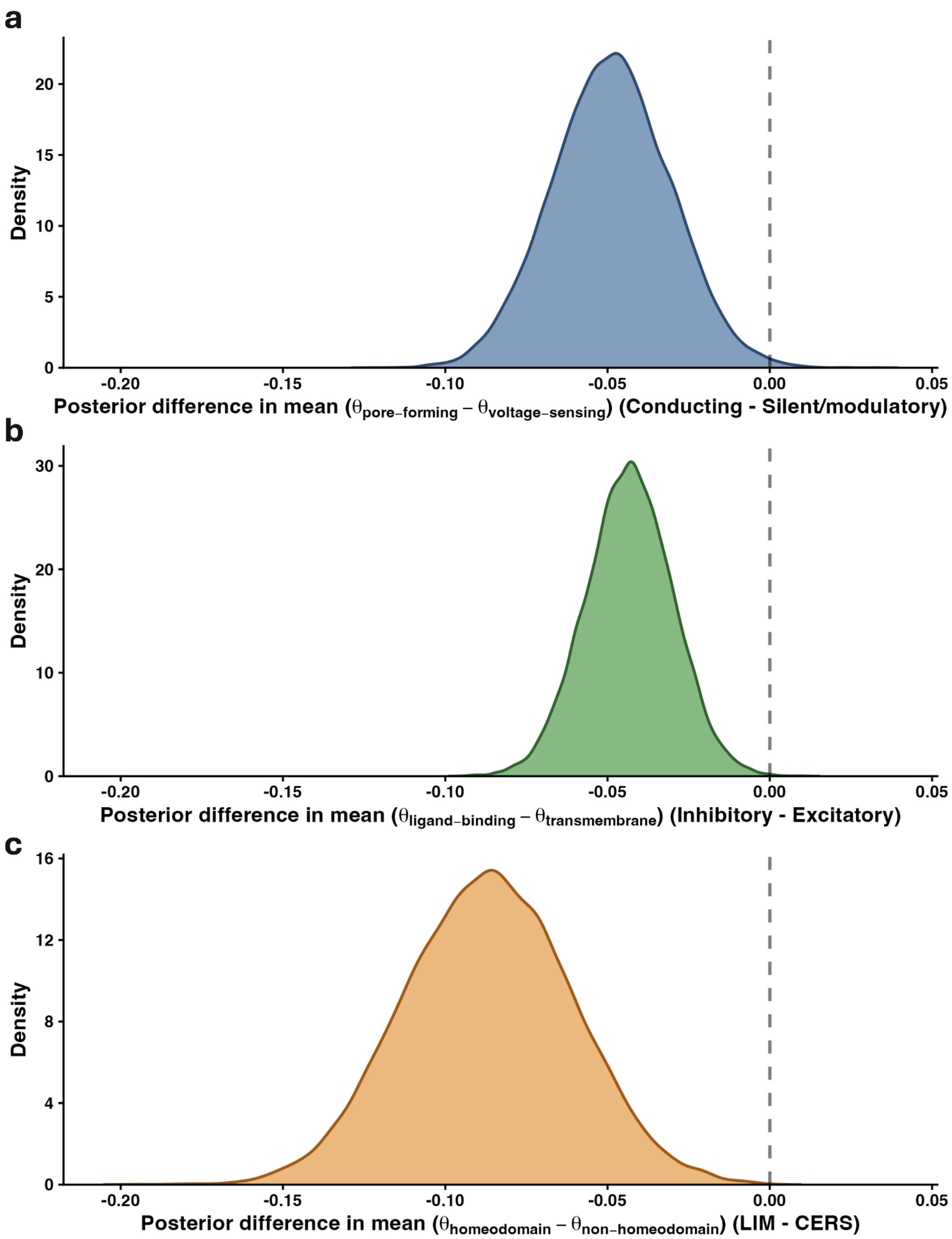


**Supplementary Figure 6: Posterior Distributions of the Difference in Means Between Gene Family Subgroups.**

**a,** Conducting Kv channels display a stronger difference in intolerance between the pore-forming and voltage-sensing domains relative to silent/modulatory Kv channels, with 99.63% of computed posterior samples below zero. **b,** Inhibitory cys-loop receptors display more intolerance on average in their ligand-binding domain than their transmembrane domain compared to excitatory receptors, with 99.94% of computed posterior samples below zero. **c,** LIM homeoboxes display more localized intolerance on average in their DNA-binding homeodomains compared to CERS homeoboxes, with 99.99% of computed posterior samples below zero.

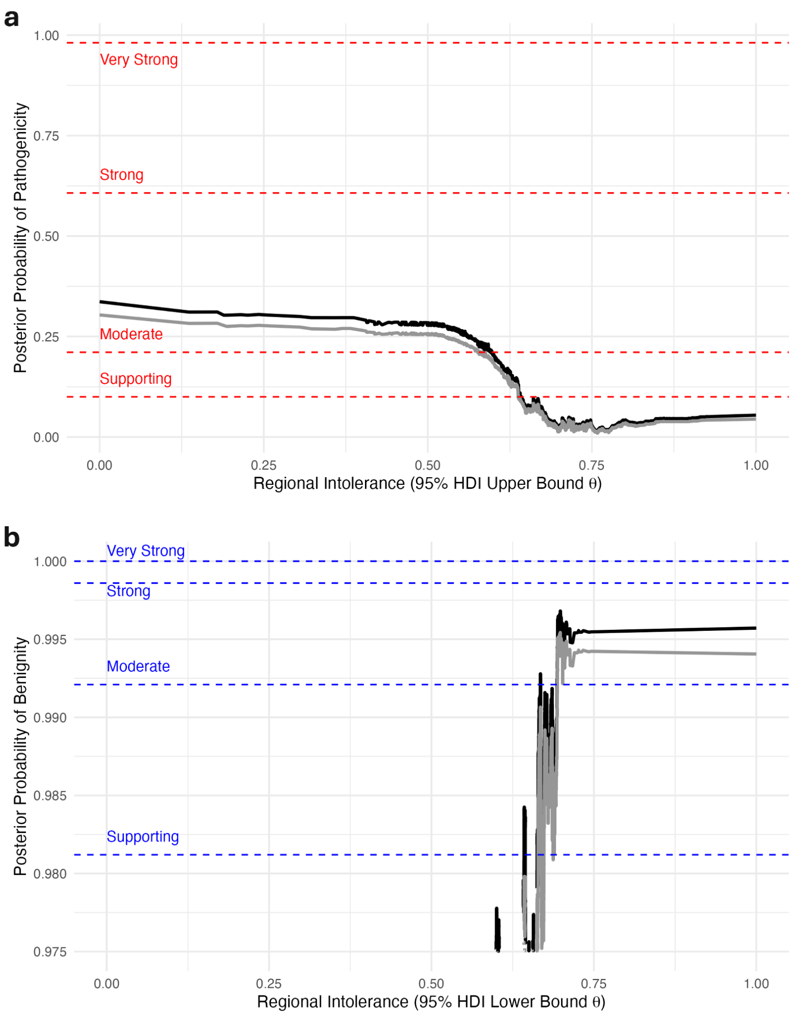


**Supplementary Figure 7: Calibration of PRIME Estimates for Pathogenicity and Benignity.**
**a**, Local posterior probabilities of pathogenicity given the upper bound of the 95% HDI of θ. This estimate is conservative in the direction of intolerance. **b**, Local posterior probabilities of benignity given the lower bound of the 95% HDI of θ. This estimate is conservative in the direction of tolerance.

| Evidence | PRIME summary statistic | Supporting | Moderate |
|---|---|---|---|
| Pathogenic | $\theta_{HDIupper}$ | (0.605139, 0.645456] | ≤ 0.605139 |
| Benign | $\theta_{HDIlower}$ | [0.670441, 0.693056) | ≥ 0.693056 |

**Supplementary Table 2: Calibrated PRIME Thresholds of Variant Pathogenicity and Benignity.** Supporting and moderate evidence thresholds of pathogenicity and benignity determined by 10,000 bootstrap iterations. "(" and ")" denote exclusion of the end value and "[" and "]" denote inclusion.

| Predictor | Strength | Predictor only | + θ supporting | + θ moderate |
|---|---|---|---|---|
| BayesDel | Supporting | 3.23 | 7.14 | 20.95 |
| BayesDel | Moderate | 8.48 | 17.14 | 49.77 |
| BayesDel | Strong | 51.73 | 107.1 | 143.19 |
| CADD | Supporting | 3.39 | 7.73 | 31.93 |
| CADD | Moderate | 6.03 | 18.79 | 48.12 |
| EA | Supporting | 3.87 | 13.13 | 58.75 |
| EA | Moderate | 12.02 | 195.83 | 99.64 |
| FATHMM | Supporting | 3.64 | 7.18 | 17.88 |
| FATHMM | Moderate | 14.5 | 24.77 | 36.38 |
| MPC | Supporting | 2.79 | 3 | 4.63 |
| MPC | Moderate | 9.59 | 8.77 | 22.21 |
| MutPred2 | Supporting | 3.92 | 4.16 | 11.18 |
| MutPred2 | Moderate | 9.58 | 16.73 | 28.26 |
| MutPred2 | Strong | 44.88 | 59.38 | 62.79 |
| PhyloP | Supporting | 4.11 | 7.27 | 27.8 |
| PhyloP | Moderate | 5.69 | 8.87 | 15.49 |
| PolyPhen2 | Supporting | 2.85 | 10.22 | 34.66 |
| PolyPhen2 | Moderate | 9.76 | 28.18 | 174.69 |
| PrimateAI | Supporting | 3.4 | 4.11 | 14.9 |
| PrimateAI | Moderate | 8.19 | 12.57 | 46.23 |
| REVEL | Supporting | 3.77 | 5.19 | 20.38 |
| REVEL | Moderate | 9.72 | 21.15 | 45.91 |
| REVEL | Strong | 52.81 | 102.14 | 139.29 |
| SIFT | Supporting | 3.31 | 7.48 | 54.2 |
| SIFT | Moderate | 6.04 | 23.23 | 30.51 |
| VEST4 | Supporting | 3.31 | 5.75 | 20.69 |
| VEST4 | Moderate | 9.56 | 16.93 | 74.67 |
| VEST4 | Strong | 35.75 | 100.63 | 103.02 |

**Supplementary Table 3: Pathogenicity Likelihood Ratios in ClinVar 2020 Dataset.**
Interval-based positive likelihood ratios for the pathogenic predictor intervals alone, predictor intervals further stratified by $\theta_{HDIupper}$ supporting evidence, and predictor intervals further stratified by $\theta_{HDIupper}$ moderate evidence. Higher values indicate higher probability of pathogenicity.

| Predictor | Strength | Predictor only | + θ supporting | + θ moderate |
|---|---|---|---|---|
| BayesDel | Supporting | 0.23 | 0.23 | 0.05 |
| BayesDel | Moderate | 0.06 | 0.06 | 0.03 |
| CADD | Supporting | 0.22 | 0.14 | 0.05 |
| CADD | Moderate | 0.06 | 0.05 | 0.02 |
| CADD | Strong | 0 | 0 | 0 |
| EA | Supporting | 0.17 | 0.18 | 0.06 |
| EA | Moderate | 0.06 | 0.05 | 0.03 |
| FATHMM | Supporting | 0.15 | 0 | 0 |
| FATHMM | Moderate | 0.05 | 0 | 0.07 |
| GERP | Supporting | 0.16 | 0.12 | 0.02 |
| GERP | Moderate | 0.12 | 0.06 | 0 |
| MutPred2 | Supporting | 0.21 | 0.27 | 0.14 |
| MutPred2 | Moderate | 0.07 | 0.09 | 0.03 |
| MutPred2 | Strong | 0 | 0 | 0 |
| PhyloP | Supporting | 0.21 | 0.17 | 0.03 |
| PhyloP | Moderate | 0.09 | 0.04 | 0.02 |
| PolyPhen2 | Supporting | 0.2 | 0.2 | 0.02 |
| PolyPhen2 | Moderate | 0.1 | 0.06 | 0.01 |
| PrimateAI | Supporting | 0.21 | 0.34 | 0.06 |
| PrimateAI | Moderate | 0.07 | 0.11 | 0.03 |
| REVEL | Supporting | 0.24 | 0.3 | 0.06 |
| REVEL | Moderate | 0.06 | 0.06 | 0.03 |
| REVEL | Strong | 0 | 0 | 0 |
| REVEL | Very strong | 0 | 0 | 0 |
| SIFT | Supporting | 0.2 | 0.26 | 0.07 |
| SIFT | Moderate | 0.08 | 0.03 | 0.04 |
| VEST4 | Supporting | 0.16 | 0.22 | 0.08 |
| VEST4 | Moderate | 0.06 | 0.1 | 0.02 |

**Supplementary Table 4: Benignity Likelihood Ratios in ClinVar 2020 Dataset.**
Interval-based positive likelihood ratios for the benign predictor intervals alone, predictor intervals further stratified by $\theta_{HDIupper}$ supporting evidence, and predictor intervals further stratified by $\theta_{HDIupper}$ moderate evidence. Lower values indicate lower probability of pathogenicity, and thus higher probability of benignity.

| Predictor | Strength | Predictor only coverage | + θ supporting coverage | + θ moderate coverage | Predictor only counts | + θ supporting counts | + θ moderate counts | Total counts |
|---|---|---|---|---|---|---|---|---|
| BayesDel | Supporting | 7.9% | 0.7% | 1.1% | 381 path / 275 benign | 46 path / 15 benign | 81 path / 9 benign | 2498 path / 5816 benign |
| BayesDel | Moderate | 11.6% | 1.1% | 2.2% | 758 path / 208 benign | 81 path / 11 benign | 171 path / 8 benign | 2498 path / 5816 benign |
| BayesDel | Strong | 8.9% | 1.1% | 1.5% | 711 path / 32 benign | 92 path / 2 benign | 123 path / 2 benign | 2498 path / 5816 benign |
| CADD | Supporting | 17.1% | 1.5% | 2.5% | 842 path / 579 benign | 93 path / 28 benign | 192 path / 14 benign | 2498 path / 5816 benign |
| CADD | Moderate | 16.1% | 1.5% | 2.3% | 966 path / 373 benign | 113 path / 14 benign | 186 path / 9 benign | 2498 path / 5816 benign |
| EA | Supporting | 9.8% | 0.9% | 1.4% | 465 path / 277 benign | 57 path / 10 benign | 102 path / 4 benign | 2284 path / 5262 benign |
| EA | Moderate | 14.1% | 1.1% | 2.3% | 892 path / 171 benign | 85 path / 1 benign | 173 path / 4 benign | 2284 path / 5262 benign |
| FATHMM | Supporting | 5.6% | 0.6% | 1.2% | 249 path / 159 benign | 34 path / 11 benign | 77 path / 10 benign | 2186 path / 5076 benign |
| FATHMM | Moderate | 7.4% | 1.0% | 0.7% | 462 path / 74 benign | 64 path / 6 benign | 47 path / 3 benign | 2186 path / 5076 benign |
| MPC | Supporting | 7.7% | 1.0% | 0.9% | 266 path / 209 benign | 37 path / 27 benign | 38 path / 18 benign | 1934 path / 4240 benign |
| MPC | Moderate | 14.0% | 2.6% | 5.6% | 704 path / 161 benign | 128 path / 32 benign | 314 path / 31 benign | 1934 path / 4240 benign |
| MutPred2 | Supporting | 7.0% | 0.7% | 0.8% | 366 path / 216 benign | 36 path / 20 benign | 58 path / 12 benign | 2495 path / 5772 benign |
| MutPred2 | Moderate | 11.8% | 1.3% | 2.2% | 787 path / 190 benign | 94 path / 13 benign | 171 path / 14 benign | 2495 path / 5772 benign |
| MutPred2 | Strong | 8.6% | 1.0% | 2.4% | 679 path / 35 benign | 77 path / 3 benign | 190 path / 7 benign | 2495 path / 5772 benign |
| PhyloP | Supporting | 23.8% | 2.6% | 4.1% | 1237 path / 706 benign | 158 path / 51 benign | 308 path / 26 benign | 2443 path / 5734 benign |
| PhyloP | Moderate | 4.1% | 0.5% | 0.9% | 235 path / 97 benign | 34 path / 9 benign | 66 path / 10 benign | 2443 path / 5734 benign |
| PolyPhen2 | Supporting | 14.2% | 1.2% | 1.8% | 564 path / 446 benign | 68 path / 15 benign | 123 path / 8 benign | 2181 path / 4916 benign |
| PolyPhen2 | Moderate | 13.1% | 1.1% | 2.2% | 758 path / 175 benign | 75 path / 6 benign | 155 path / 2 benign | 2181 path / 4916 benign |
| PrimateAI | Supporting | 12.8% | 1.5% | 2.0% | 594 path / 408 benign | 74 path / 42 benign | 134 path / 21 benign | 2355 path / 5499 benign |
| PrimateAI | Moderate | 12.5% | 2.1% | 4.0% | 765 path / 218 benign | 140 path / 26 benign | 297 path / 15 benign | 2355 path / 5499 benign |
| REVEL | Supporting | 6.8% | 0.7% | 1.1% | 343 path / 211 benign | 38 path / 17 benign | 79 path / 9 benign | 2442 path / 5669 benign |
| REVEL | Moderate | 11.6% | 1.1% | 2.3% | 762 path / 182 benign | 82 path / 9 benign | 178 path / 9 benign | 2442 path / 5669 benign |
| REVEL | Strong | 8.2% | 1.1% | 1.5% | 637 path / 28 benign | 88 path / 2 benign | 120 path / 2 benign | 2442 path / 5669 benign |
| SIFT | Supporting | 8.0% | 0.5% | 1.0% | 333 path / 234 benign | 29 path / 9 benign | 70 path / 3 benign | 2128 path / 4943 benign |
| SIFT | Moderate | 20.5% | 1.9% | 3.0% | 1048 path / 403 benign | 120 path / 12 benign | 197 path / 15 benign | 2128 path / 4943 benign |
| VEST4 | Supporting | 9.4% | 0.9% | 1.4% | 426 path / 308 benign | 48 path / 20 benign | 95 path / 11 benign | 2291 path / 5489 benign |
| VEST4 | Moderate | 14.6% | 1.6% | 2.5% | 906 path / 227 benign | 106 path / 15 benign | 187 path / 6 benign | 2291 path / 5489 benign |
| VEST4 | Strong | 5.1% | 0.6% | 0.6% | 373 path / 25 benign | 42 path / 1 benign | 43 path / 1 benign | 2291 path / 5489 benign |

**Supplementary Table 5: Pathogenicity Threshold Coverage of ClinVar 2020 Variants.**
Coverage of total variants in the dataset for the pathogenic predictor intervals alone, predictor intervals further stratified by $\theta_{HDIupper}$ supporting evidence, and predictor intervals further stratified by $\theta_{HDIupper}$ moderate evidence.

| Predictor | Strength | Predictor only coverage | + θ supporting coverage | + θ moderate coverage | Predictor only counts | + θ supporting counts | + θ moderate counts | Total counts |
|---|---|---|---|---|---|---|---|---|
| BayesDel | Supporting | 17.8% | 1.8% | 1.1% | 131 path / 1351 benign | 13 path / 134 benign | 2 path / 88 benign | 2498 path / 5816 benign |
| BayesDel | Moderate | 32.0% | 4.3% | 3.8% | 63 path / 2594 benign | 9 path / 349 benign | 4 path / 312 benign | 2498 path / 5816 benign |
| CADD | Supporting | 18.6% | 1.7% | 1.2% | 131 path / 1418 benign | 8 path / 134 benign | 2 path / 101 benign | 2498 path / 5816 benign |
| CADD | Moderate | 23.9% | 2.9% | 2.8% | 47 path / 1938 benign | 5 path / 237 benign | 2 path / 232 benign | 2498 path / 5816 benign |
| CADD | Strong | 2.6% | 0.4% | 0.5% | 0 path / 217 benign | 0 path / 34 benign | 0 path / 43 benign | 2498 path / 5816 benign |
| EA | Supporting | 25.5% | 2.5% | 2.1% | 130 path / 1791 benign | 14 path / 177 benign | 4 path / 151 benign | 2284 path / 5262 benign |
| EA | Moderate | 13.6% | 1.8% | 1.2% | 28 path / 999 benign | 3 path / 130 benign | 1 path / 87 benign | 2284 path / 5262 benign |
| FATHMM | Supporting | 2.3% | 0.4% | 0.6% | 10 path / 160 benign | 0 path / 27 benign | 0 path / 42 benign | 2186 path / 5076 benign |
| FATHMM | Moderate | 0.7% | 0.1% | 0.5% | 1 path / 49 benign | 0 path / 4 benign | 1 path / 35 benign | 2186 path / 5076 benign |
| GERP | Supporting | 22.3% | 2.3% | 2.6% | 119 path / 1702 benign | 9 path / 176 benign | 2 path / 212 benign | 2442 path / 5730 benign |
| GERP | Moderate | 3.8% | 0.5% | 0.5% | 15 path / 298 benign | 1 path / 42 benign | 0 path / 40 benign | 2442 path / 5730 benign |
| MutPred2 | Supporting | 17.4% | 1.8% | 1.1% | 120 path / 1316 benign | 16 path / 135 benign | 5 path / 84 benign | 2495 path / 5772 benign |
| MutPred2 | Moderate | 34.1% | 4.3% | 3.9% | 86 path / 2731 benign | 14 path / 343 benign | 4 path / 321 benign | 2495 path / 5772 benign |
| MutPred2 | Strong | 0.1% | 0.0% | 0.0% | 0 path / 8 benign | 0 path / 3 benign | 0 path / 0 benign | 2495 path / 5772 benign |
| PhyloP | Supporting | 22.9% | 2.7% | 2.2% | 156 path / 1717 benign | 15 path / 203 benign | 2 path / 179 benign | 2443 path / 5734 benign |
| PhyloP | Moderate | 11.1% | 1.4% | 1.5% | 35 path / 874 benign | 2 path / 110 benign | 1 path / 125 benign | 2443 path / 5734 benign |
| PolyPhen2 | Supporting | 15.4% | 1.6% | 1.5% | 90 path / 1004 benign | 9 path / 103 benign | 1 path / 102 benign | 2181 path / 4916 benign |
| PolyPhen2 | Moderate | 22.9% | 2.6% | 2.3% | 66 path / 1558 benign | 5 path / 176 benign | 1 path / 162 benign | 2181 path / 4916 benign |
| PrimateAI | Supporting | 14.5% | 1.8% | 1.1% | 94 path / 1047 benign | 18 path / 122 benign | 2 path / 84 benign | 2355 path / 5499 benign |
| PrimateAI | Moderate | 20.9% | 3.3% | 3.6% | 47 path / 1594 benign | 12 path / 247 benign | 4 path / 280 benign | 2355 path / 5499 benign |
| REVEL | Supporting | 13.8% | 1.3% | 0.9% | 105 path / 1011 benign | 12 path / 94 benign | 2 path / 72 benign | 2442 path / 5669 benign |
| REVEL | Moderate | 32.8% | 4.1% | 3.8% | 70 path / 2590 benign | 9 path / 325 benign | 4 path / 307 benign | 2442 path / 5669 benign |
| REVEL | Strong | 1.6% | 0.3% | 0.3% | 0 path / 132 benign | 0 path / 25 benign | 0 path / 25 benign | 2442 path / 5669 benign |
| REVEL | Very strong | 0.1% | 0.0% | 0.1% | 0 path / 12 benign | 0 path / 4 benign | 0 path / 5 benign | 2442 path / 5669 benign |
| SIFT | Supporting | 20.3% | 2.1% | 1.5% | 113 path / 1322 benign | 15 path / 134 benign | 3 path / 102 benign | 2128 path / 4943 benign |
| SIFT | Moderate | 18.6% | 2.1% | 1.6% | 43 path / 1272 benign | 2 path / 145 benign | 2 path / 112 benign | 2128 path / 4943 benign |
| VEST4 | Supporting | 12.6% | 1.2% | 0.8% | 63 path / 918 benign | 8 path / 89 benign | 2 path / 58 benign | 2291 path / 5489 benign |
| VEST4 | Moderate | 36.0% | 4.4% | 3.9% | 63 path / 2739 benign | 14 path / 332 benign | 3 path / 303 benign | 2291 path / 5489 benign |

**Supplementary Table 6: Benignity Threshold Coverage of ClinVar 2020 Variants.**
Coverage of total variants in the dataset for the benign predictor intervals alone, predictor intervals further stratified by $\theta_{HDIlower}$ supporting evidence, and predictor intervals further stratified by $\theta_{HDIlower}$ moderate evidence.

| Gene Set | Autosomal Dominant | Autosomal Recessive | Dominant-Negative | Haploinsufficient | Gain-of-Function | De Novo |
|---|---|---|---|---|---|---|
| **Autosomal Dominant** | 1606 (100.0%) | 459 (28.6%) | 547 (34.1%) | 225 (14.0%) | 406 (25.3%) | 989 (61.6%) |
| **Autosomal Recessive** | 459 (16.2%) | 2830 (100.0%) | 283 (10.0%) | 79 (2.8%) | 166 (5.9%) | 406 (14.3%) |
| **Dominant-Negative** | 547 (80.9%) | 283 (41.9%) | 676 (100.0%) | 100 (14.8%) | 183 (27.1%) | 429 (63.5%) |
| **Haploinsufficient** | 225 (81.2%) | 79 (28.5%) | 100 (36.1%) | 277 (100.0%) | 63 (22.7%) | 192 (69.3%) |
| **Gain-of-Function** | 406 (83.5%) | 166 (34.2%) | 183 (37.7%) | 63 (13.0%) | 486 (100.0%) | 321 (66.0%) |
| **De Novo** | 989 (75.8%) | 406 (31.1%) | 429 (32.9%) | 192 (14.7%) | 321 (24.6%) | 1305 (100.0%) |

**Supplementary Table 7: OMIM Gene Set Overlap.**

Overlap of gene counts within derived OMIM disease gene sets. Percentages are computed for the gene set in the corresponding row. Genes may have both dominant and recessive phenotypes in OMIM, explaining the overlap of gene sets with dominant effects with the autosomal recessive group.
